# Mendelian randomization accounting for correlated and uncorrelated pleiotropic effects using genome-wide summary statistics

**DOI:** 10.1101/682237

**Authors:** Jean Morrison, Nicholas Knoblauch, Joseph Marcus, Matthew Stephens, Xin He

**Affiliations:** Department of Human Genetics, University of Chicago; Department of Statistics, University of Chicago

## Abstract

Mendelian randomization (MR) is a valuable tool for detecting evidence of causal relationships using genetic variant associations. Opportunities to apply MR are growing rapidly with the number of genome-wide association studies (GWAS) with publicly available results. However, existing MR methods rely on strong assumptions that are often violated, leading to false positives. Many methods have been proposed loosening these assumptions. However, it has remained challenging to account for correlated pleiotropy, which arises when variants affect both traits through a heritable shared factor. We propose a new MR method, Causal Analysis Using Summary Effect Estimates (CAUSE), that accounts for correlated and uncorrelated horizontal pleiotropic effects. We demonstrate in simulations that CAUSE is more robust to correlated pleiotropy than other methods. Applied to traits studied in recent GWAS, we find that CAUSE detects causal relationships with strong literature support and avoids identifying most unlikely relationships. Our results suggest that many pairs of traits identified as causal using alternative methods may be false positives due to horizontal pleiotropy.

## 1 Introduction

Inferring causal relationships between traits is important for understanding the etiology of disease and designing new treatments. Randomized trials are considered the gold standard for inferring causality, but are expensive and sometimes impossible. Observational studies are often a feasible alternative, but observational associations may be biased by confounding and reverse causality.

Mendelian randomization (MR) is a potentially powerful approach to studying causal relationships using data from observational genetic studies. The key idea of MR is to treat genotypes as naturally occurring “randomizations” [1, 2, 3, 4]. Suppose we are interested in the causal effect, *γ*, of *M* (for “Mediator”) on *Y*. If a variant *G*_*j*_ affects *M* and doesn’t affect *Y* through pathways not mediated by *M* (see Figure 1), then *G*_*j*_ can be viewed as an un-confounded proxy for *M*. Under these assumptions

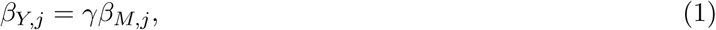

where *β*_*Y,j*_ is the association of *G*_*j*_ with *Y, β*_*M,j*_ is the association of *G*_*j*_ with *M*. This relationship is the core of simple MR methods. Many methods based on Equation (1), including the commonly used inverse variance weighted (IVW) regression, first obtain estimates of *β*_*Y,j*_ and *β*_*M,j*_ for several variants strongly associated with *M*, and then estimate *γ* by regressing the estimates of *β*_*Y,j*_ on the estimates of *β*_*M,j*_ [5].

**Figure 1:**
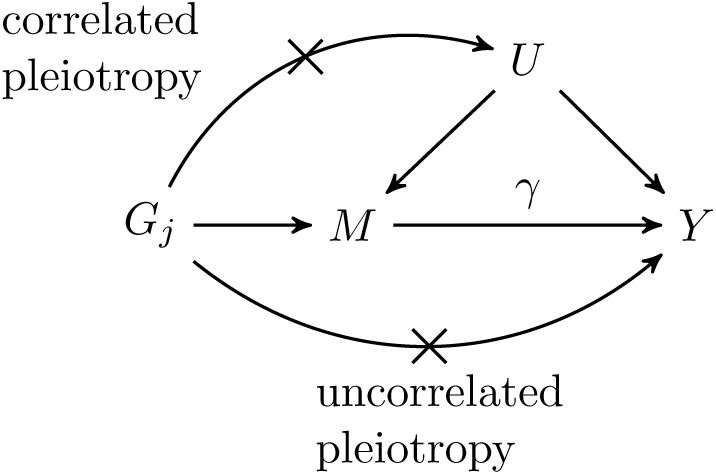
Causal diagram assumed by simple MR. Arrows indicate causal effects. Crosses mark causal effects that are assumed absent by simple MR methods. The causal effect of trait *M* on trait *Y, γ*, is the target of inference. Variant *G*_*j*_ affects trait *M* but is assumed to have no effects on *Y* that are not mediated through *M*. Pleiotropic effects on *Y* that are independent of the effect of *G*_*j*_ on *M* are referred to as uncorrelated pleiotropy. Effects that are mediated by a shared factor, *U*, are referred to as correlated pleiotropy.

A variant that affects *M* and also affects *Y* through a pathway not mediated by a causal effect of *M* on *Y* is termed *horizontally pleiotropic*. Historically, horizontal pleiotropy was assumed to be rare because it was thought that each variant had few biological effects and that most confounders (*U* in Figures 1) were non-heritable environmental risks. It has subsequently become clear that horizontal pleioptropy is common and leads to MR false positives if it is not accounted for [6, 7]. Accounting for these effects has been a major focus of ongoing MR research [4]. It is helpful to distinguish between two types of horizontal pleiotropy: *uncorrelated pleiotropy*, where horizontal pleiotropic effects on *Y* are uncorrelated with effects on *M*, and *correlated pleiotropy*, where horizontal pleiotropic effects on *Y* are correlated with effects on *M*. Uncorrelated pleiotropy occurs when a genetic variant affects *Y* and *M* through separate mechanisms, whereas correlated pleiotropy occurs when a genetic variant affects *Y* and *M* through a shared heritable factor, *U* (Figure 1). *U* may represent a shared biological process or pathway. Both types of pleiotropy may occur for any pair of traits.

Of the two types, uncorrelated horizontal pleiotropy is easier to account for. If it is mean zero, it only adds noise to the relationship in Equation (1) rather than inducing a systematic correlation between *β*_*Y,j*_ and *β*_*M,j*_. Several methods have recently been developed to deal with this, including Egger regression [8, 9] which allows for directional, or non-mean zero uncorrelated pleiotropy, and several methods that rely on outlier removal including GSMR [10] and MR-PRESSO [7]. However, because these methods do not model correlated pleiotropy, they remain vulnerable to false positives in its presence. Genetic correlation has been found between many pairs of traits including some that are unlikely to be causally linked, which suggests that correlated pleiotropy is common and may be an important source of false positives in MR analyses [11, 12, 13, 14]. If a potential shared factor is suspected and measured, false positives can be reduced by eliminating variants associated with the shared factor or adjusting for it using multivariable MR [15]. Two methods, the weighted median approach [16] and the weighted mode approach [17], allow for some correlated pleiotropy. These methods rely respectively on the assumption that fewer than 50% of variants display any form of horizontal pleiotropy and that mode of pleiotropic effects is zero. Another method, LCV [14], does not directly test for a causal effect, but estimates the “genetic causality proportion”, with higher magnitude estimates suggestive of a causal effect and lower magnitude estimates suggestive of correlated pleiotropy.

We present a new MR method that accounts for both uncorrelated and correlated pleiotropy. Causal Analysis Using Summary Effect Estimates (CAUSE) uses genome-wide summary statistics to identify pairs of traits that are consistent with a causal effect. The intuition behind CAUSE is that a causal effect of *M* on *Y* leads to correlation between *β*_*M,j*_ and *β*_*Y,j*_ for all variants with non-zero effect on *M* (Figure 2b). On the other hand, if *M* has no causal effect on *Y*, but both traits are affected by a shared factor, then only the subset of *M* effect variants acting on the shared factor will have correlated estimates (Figure 2a). CAUSE uses this distinction to differentiate causal effects from correlated pleiotropy. In addition to accounting for correlated pleiotropy, CAUSE differs from other MR methods by using information from all variants, rather than only those most strongly associated with *M*. This can improve power when the GWAS for trait *M* is under-powered. CAUSE provides a test-statistic, an estimate of the causal effect, and a variant-level summary indicating how each variant contributes to the overall test, and which are likely to be acting through a shared factor.

**Figure 2:**
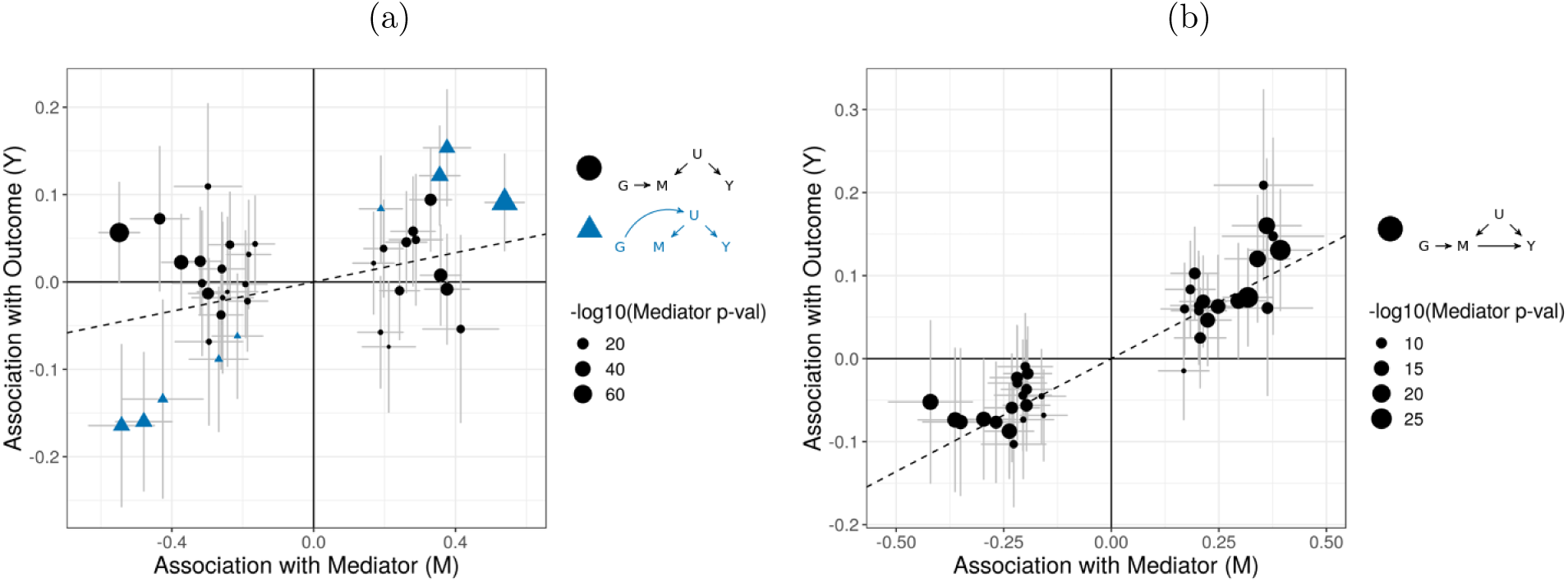
Simulated data illustrating (a) the pattern induced by a shared factor (correlated pleiotropy) without a causal effect; and (b) the pattern induced by a causal effect. In both plots, effect size estimates are indicated by points. Error bars around points have length 1.96 times the standard error on each side. Only variants that are strongly associated with *M* (*p* < 5 · 10^−8^) are shown. (a) Summary statistics are simulated for two traits with no causal effect. 15% of the *M* effect variants act on *M* through a shared factor that affects both *M* and *Y* (triangles). Although there are many variants associated with *M* and not with *Y*, the IVW estimator (dotted line) is significantly different from zero (*p* = 0.01), driven by the effect size correlation of the 15% pleiotropic variants (triangles). (b) Summary statistics are simulated under a model where *M* causally affects *Y*. Here *every* variant associated with *M* is also associated with *Y* and the effect sizes are correlated.

We demonstrate in simulations that CAUSE makes fewer false detections in the presence of correlated pleiotropy than existing methods. In applications to GWAS data, CAUSE identifies plausible causal relationships while avoiding many likely false positives. CAUSE fills a gap in existing methodology by accounting for correlated pleiotropy created by unknown or unmeasured shared factors.

## 2 Results

### 2.1 CAUSE models uncorrelated and correlated pleiotropy

Using CAUSE, we assess whether genome-wide variant effect estimates and standard errors for *M* and *Y* are consistent with a causal effect. In a simple MR analysis, variants with strong associations with *M* are selected and assumed to follow the causal diagram in Figure 1. In our proposal, illustrated in Figure 3, we use all variants, but assume that most have no effect on either trait. We assume that the majority of variants that do affect *M* follow the causal diagram in Figure 3a, which is the same as the diagram in Figure 1 with the addition of an uncorrelated pleiotropic effect. We allow a small proportion, *q*, of *M* effect variants to follow the causal diagram in Figure 3b, acting on both *M* and *Y* through an unobserved heritable shared factor, *U*. Under this model, the relationship between *β*_*Y,j*_ and *β*_*M,j*_ is

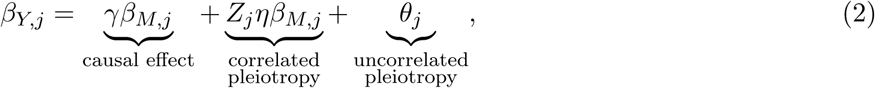

where *Z*_*j*_ is an indicator that is 1 if *G*_*j*_ acts on *M* through *U* and 0 otherwise, and *η* is the effect of *U* on *Y*. This relationship is an extension of Equation (1) that includes terms allowing for both types of horizontal pleiotropic effects. We assume that *q* is small, so *Z*_*j*_ is equal to 0 for most variants (see Methods).

**Figure 3:**
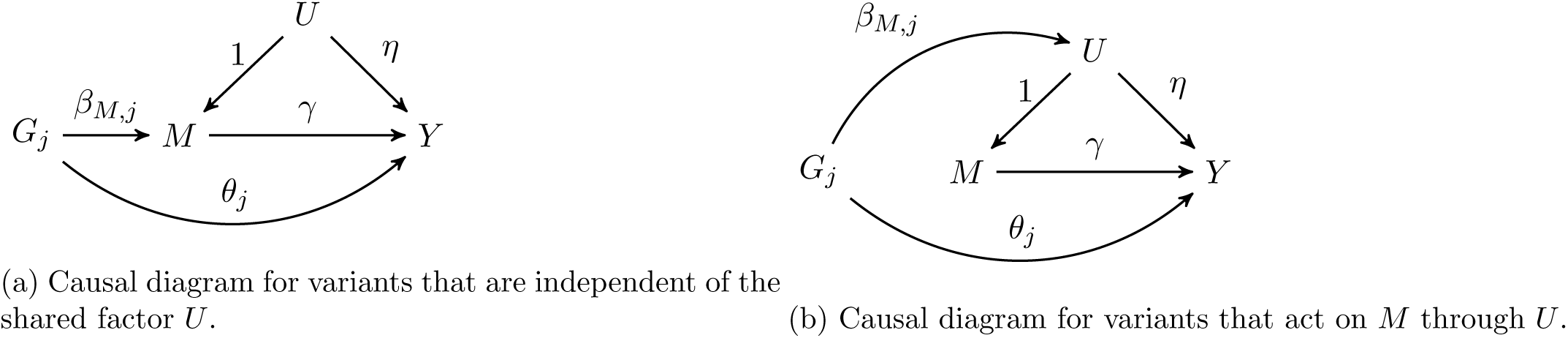
CAUSE assumes that variants affect trait *M* through one of two mechanisms. A proportion 1 − *q* of variants have the causal diagram in (a), while the remaining proportion, *q*, have the causal diagram in (b). (a) *G*_*j*_ has an effect on *M* of size *β*_*M,j*_ and is independent of *U*. *G*_*j*_ can affect *Y* through two pathways: through the uncorrelated effect, *θ*_*j*_, and through the effect of *M* on *Y* (if *γ* ≠ 0). (b) The effect of *G*_*j*_ on *M* is mediated by *U*. We have assumed that *U* is scaled so that its effect on *M* is 1. This makes the effect of *G*_*j*_ on *U* equal to the effect of *G*_*j*_ on *M, β*_*M,j*_. The effect of *U* on *Y* is equal to *η*, an unknown parameter. *G*_*j*_ can affect *Y* through three pathways: through *θ*_*j*_, through the causal effect of *M* on *Y*, and through the effect of *U* on *Y*.

Equation (2) allows us to model the patterns in Figure 2. If *γ* = 0 and *q* = 0 (no causal effect and no shared factor), then *β*_*Y,j*_ and *β*_*M,j*_ are uncorrelated for all *j*. If there is a shared factor and no causal effect (*γ* = 0, *q* and *η* non-zero), then *β*_*Y,j*_ and *β*_*M,j*_ are correlated for variants with *Z*_*j*_ = 1 (Figure 2a). If there is a causal effect (*γ* ≠ 0), *β*_*Y,j*_ and *β*_*M,j*_ are correlated for all variants (Figure 2b). Including all variants allows us to model uncertainty about variant effects on *M* and gain information from weakly associated variants (see Methods). For computational simplicity, we use a likelihood for independent variants and prune variants for LD before estimating posterior distributions and computing test statistics (see Methods and Supplementary Note Section SN4).

We assess whether GWAS summary statistics for the two traits are consistent with a causal effect by comparing two nested models. The *sharing model* has *γ* fixed at 0, allowing for pleiotropic effects but no causal effect, and the *causal model* in which *γ* is a free parameter. We compare the models using the expected log pointwise posterior density (ELPD; [18]), a Bayesian model comparison approach that estimates how well the posterior distributions of a particular model are expected to predict a new set data (see Methods). We produce a one-sided *p*-value testing that the sharing model fits the data at least as well as the causal model. If this hypothesis is rejected, we conclude that the data are consistent with a causal effect.

The causal diagram in Figure 3b is related to the model used in the latent causal variable (LCV) method proposed by O’Connor and Price [14] (see Supplementary Note Section SN5). Unlike other MR methods, LCV does not provide a test for a causal effect. LCV considers the proportion of heritability of each trait that is mediated by a shared factor, summarized as the “genetic causality proportion” (GCP) which ranges from −1 to 1. A causal model has a GCP of 1 or −1, depending on the direction of effect. O’Connor and Price [14] consider a large magnitude estimated GCP more likely to be causal and use a threshold of 0.6 to identify suggestive pairs.

### 2.2 CAUSE can distinguish causality from correlated pleiotropy in simulations

We simulate summary statistics with realistic LD patterns to assess CAUSE in a variety of scenarios and compare performance with other MR methods. Both traits are simulated with a polygenic trait architecture – an average of 1,000 variants with normally distributed effects contribute to a total heritability of 0.25. The power of trait *M* and *Y* GWAS can influence performance of all methods, so we consider low power (median 8 genome-wide significant loci) and a high power (median 107 genome-wide significant loci) settings for both traits. Power is controlled by adjusting the simulated GWAS sample size and genome-wide significance is defined as *p* < 5 · 10^−8^. Variant effects on trait *Y* are generated from the relationship in Equation (2) and effect estimates are then simulated from true effects and LD structure using results of Zhu and Stephens [19] (see Methods).

We compare CAUSE to six MR methods: IVW regression using multiplicative random effects [15], Egger regression with random effects [8], GSMR [10], MR-PRESSO [7], the weighted median estimator [16], and the modal estimator [17]. IVW regression, Egger regression, GSMR and MR-PRESSO assume that no variants exhibit correlated pleiotropy. The weighted median and modal methods are more permissive, assuming respectively that fewer than half the variants are pleiotropic and that the modal pleiotropic effect is zero. All six alternative methods require selection of variants with strong evidence of association with trait *M*. We follow the typical practice of selecting variants with *p* < 5·10^−8^ for association with *M* and pruning for LD so that no pair of variants has pairwise *r*^2^ > 0.1.

We first evaluate the robustness of each method to correlated pleiotropy by simulating data with no causal effect and a proportion *q* = 0 to 0.5 of variants acting through a shared factor (Figure 4a). We consider weak 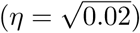 and strong 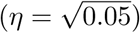 shared factor effects. CAUSE controls the false positive rate in the presence of correlated pleiotropy (*q* > 0) better than other methods except for Egger regression and the modal estimator. CAUSE makes more false detections when *q* is large, the confounder effect is large, and the GWAS have higher power. This is expected because the data pattern resulting from a high proportion of shared variants is similar to the pattern that results from a causal effect.

**Figure 4:**
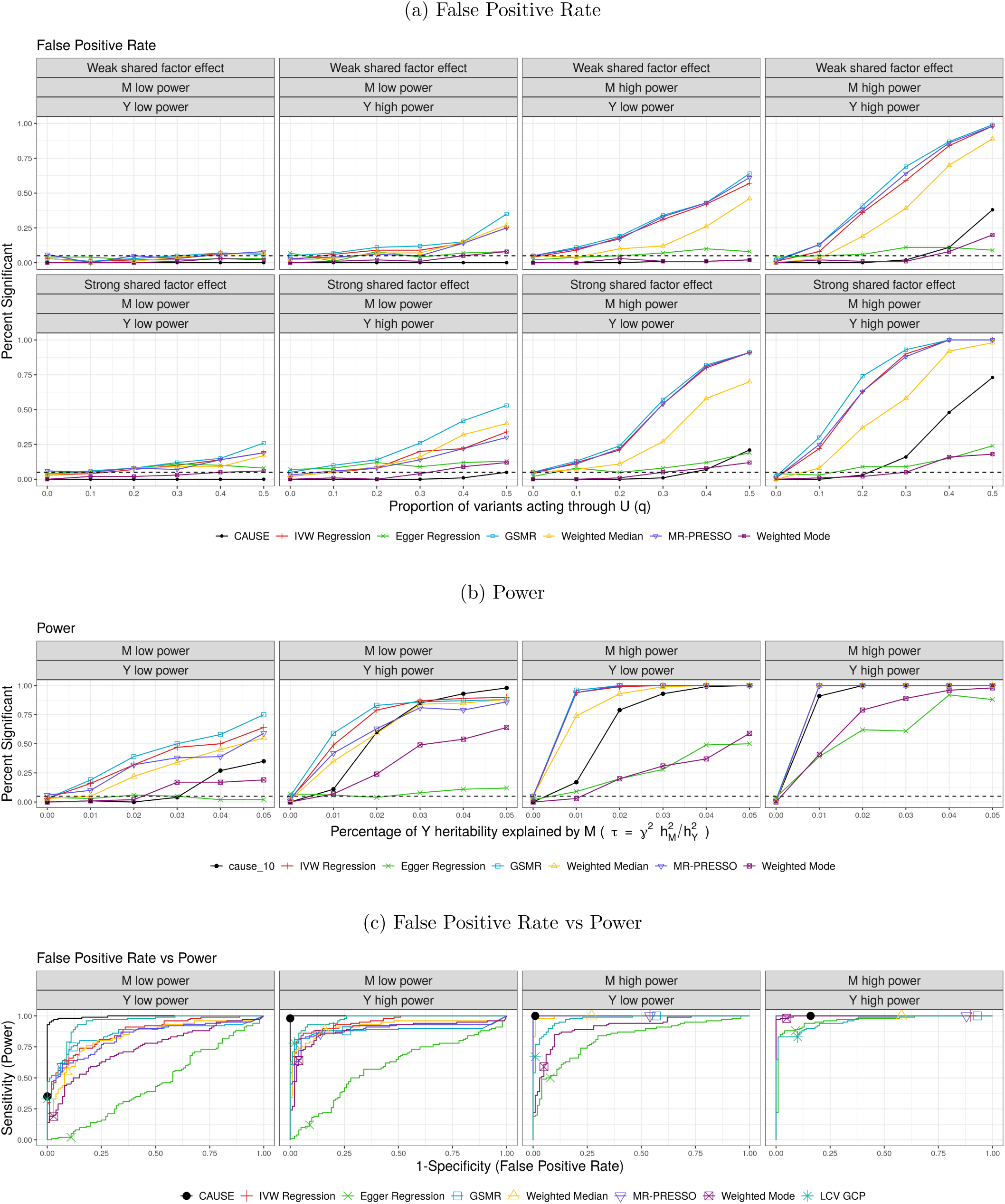
Performance of CAUSE and other MR methods in simulated data. (a) False positive rate averaged over 100 simulated data sets in settings with no causal effect and a proportion of correlated pleiotropic variants (*q*) ranging from 0 to 50%. (b) Power averaged over 100 simulated data sets in settings with a causal effect and no shared factor. (c) Comparison of false positive-power trade-off. We compare the power when 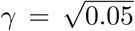 and there is no shared factor to the false positive rate when there is no causal effect, but a proportion *q* = 0.3 of variants act through a shared factor with effect 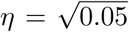 on *Y*. There are 100 simulations each in the causal and non-causal scenarios. Curves are created by varying the significance threshold. Points indicate the power and false positive rate achieved at a threshold of *p* ≤ 0.05 or *GĈP*0.06 for LCV.

We next compare the power of each method when there is a true causal effect of *M* on *Y* and no shared factor (Figure 4b). CAUSE has substantially better power than Egger regression and the modal estimator, the two methods that were able to control the false positive rate for high levels of correlated pleiotropy. CAUSE has somewhat lower power than other methods in most settings. However, when the trait *M* GWAS has low power and trait *Y* has high power, CAUSE can achieve better power than other methods for larger causal effects. This is a result of using all variants genome-wide, rather than only those reaching genome-wide significance. The posterior median of *γ* can be taken as a point estimate of the causal effect. This estimate tends to be shrunk slightly towards zero compared to alternative methods but has a substantially lower mean squared error than estimates obtained using the weighted mode or Egger regression (Supplementary Note Section SN6.2). In simulations with both a causal effect and a shared factor, CAUSE maintains similar power when *η* has a smaller magnitude than *γ* but loses power when *η* is large and has opposite sign to *γ* (Supplementary Note Section SN6.3).

Although in most settings, CAUSE has both lower power and lower false positive rate than other methods, the difference between CAUSE and other methods is not simply calibration. In Figure 4c we compare the trade-off between power and false positive rate for CAUSE and other methods, additionally including the LCV GCP estimate. CAUSE is better able to distinguish a causal scenario from a non-causal scenario with 30% correlated pleiotropy. This pattern is consistent for levels of correlated pleiotropy between 0% and 50% (Figure S2, Table S3). Additionally, CAUSE is better or equal to other methods at discriminating scenarios with a causal effect and a shared factor from those with only a shared factor (Supplementary Note Section SN6.3).

### 2.3 CAUSE avoids false positives due to reverse causality

Many MR methods obtain false positives when there is a true causal effect of *Y* on *M* but a test is performed for an effect of *M* on *Y* (reverse causal effects). This occurs because a reverse effect creates a similar pattern of variant associations to the pattern created by a shared factor. If *Y* affects *M*, some variants will have correlated effects on both traits. However, for most polygenic traits, there will be some variants that affect *M* through pathways independent of *Y* and therefore do not exhibit this correlation. Since CAUSE models correlated pleiotropy, it can avoid false positives due to reverse causality as long as there are a sufficient number of *M* effect variants that are independent of the causal path from *Y* to *M*.

To verify this expectation, we simulate data with a causal effect of *Y* on *M* and no shared factor and test for an effect of *M* on *Y* using CAUSE and other methods. In all scenarios, the expected heritability of both *M* and *Y* is 0.25. For trait *Y*, 1000 variants with normally distributed effects explain this heritability. The causal effect of *Y* on *M* explains 20% of the trait *M* heritability while the rest comes from a set of 250, 500, or 1000 variants (see Methods). CAUSE and the modal estimator both avoid false positives in all settings while the other methods do not (Figure 5). More false positives from other methods occur when the number of independent *M* effect variants is low. In these scenarios, many of the variants significantly associated with *M* act on *M* through *Y* and thus have correlated effects. Some false positives from alternative methods could be avoided by carefully filtering variants, for example using Steiger filtering [20]. This analysis demonstrates that CAUSE is able to avoid false positives from reverse effects without filtering, when there are independent variants associated with *M* independent of *Y*.

**Figure 5:**
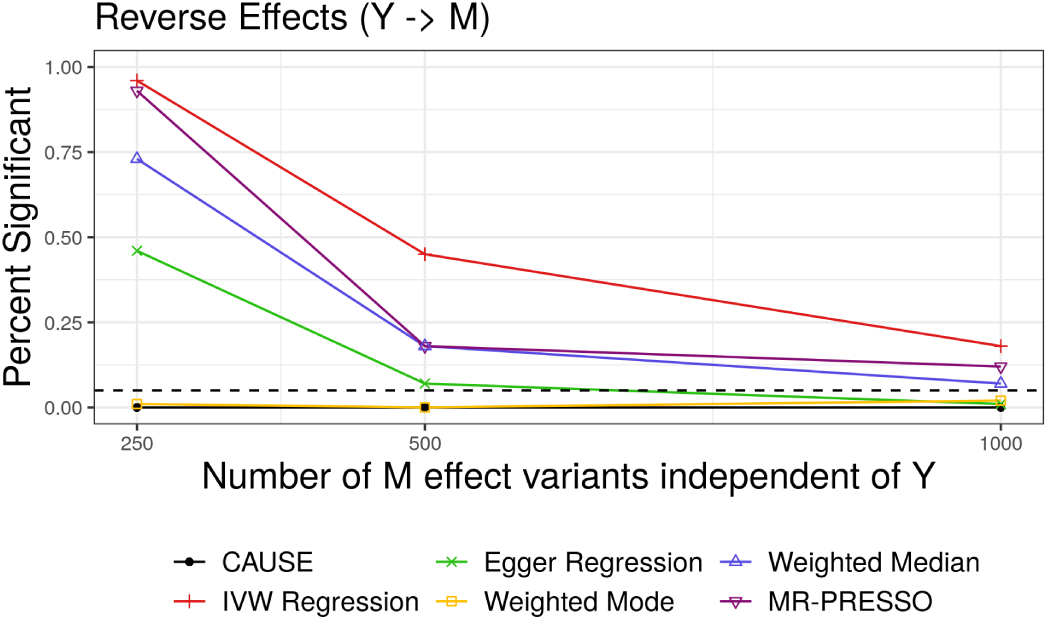
False positives resulting from reverse causality. Only CAUSE and the weighted mode control the false positive rate.

CAUSE may still find evidence of causality in both directions for some pairs of traits for several reasons. First, there may be a causal effect in one direction that accounts for nearly all of the heritability of the downstream/child trait. Second, there may be causal effects in both directions. Third, the two traits may be very closely related and share nearly all genetic variants despite having no causal relationship.

### 2.4 Identifying causal risk factors of common diseases

We use CAUSE and other methods to test for effects of twelve possible risk factors for cardio-metabolic diseases coronary artery disease (CAD), stroke, and type 2 diabetes (T2D). These include anthropometric traits, lipids, blood pressure, fasting glucose, smoking, and alcohol consumption (Table S1). In order to increase the number of tested relationships that are unlikely to be causal, we also test for effects of each mediator on asthma, though not all of these relationships are negative controls. Focusing on well studied risk factors and diseases allows us to compare results of each method with evidence from scientific literature.

Table 1 summarizes *p*-values and estimated effect direction for each method as well as GCP estimates from LCV and genetic correlation estimates from LD-score regression (see also Figure S3 and Table S5). Full lists of variants used for each MR method are provided in Tables S9 and S10. We classified trait pairs in Table 1 by evidence of causality from existing literature (see Supplementary Note Section SN7) and ordered them by strength of evidence for genetic correlation. CAUSE obtains a *p*-value less than 0.05 for 8 of 9 relationships in the considered causal category, and for 4 of 10 relationships with literature support. Of the remaining trait pairs in these two categories, 4 are not identified as causal by any method. The remaining negative results from CAUSE are discussed in Supplementary Note Section SN8. Using LCV, very few pairs of traits have estimated GCP larger than 60%, the threshold used by O’Connor and Price as suggestive of a causal relationship, however many pairs have moderate GCP estimates that are significantly different from zero. Variant estimates for smoking and CAD, a strong causal effect detected by all methods, are shown in Figure 6a colored by individual variant contribution to the CAUSE test statistic. Although there is some heterogeneity in effect size correlation, the large number of variants with correlated effects provide enough evidence for CAUSE to reject the sharing model in favor of the causal model.

**Table 1:**
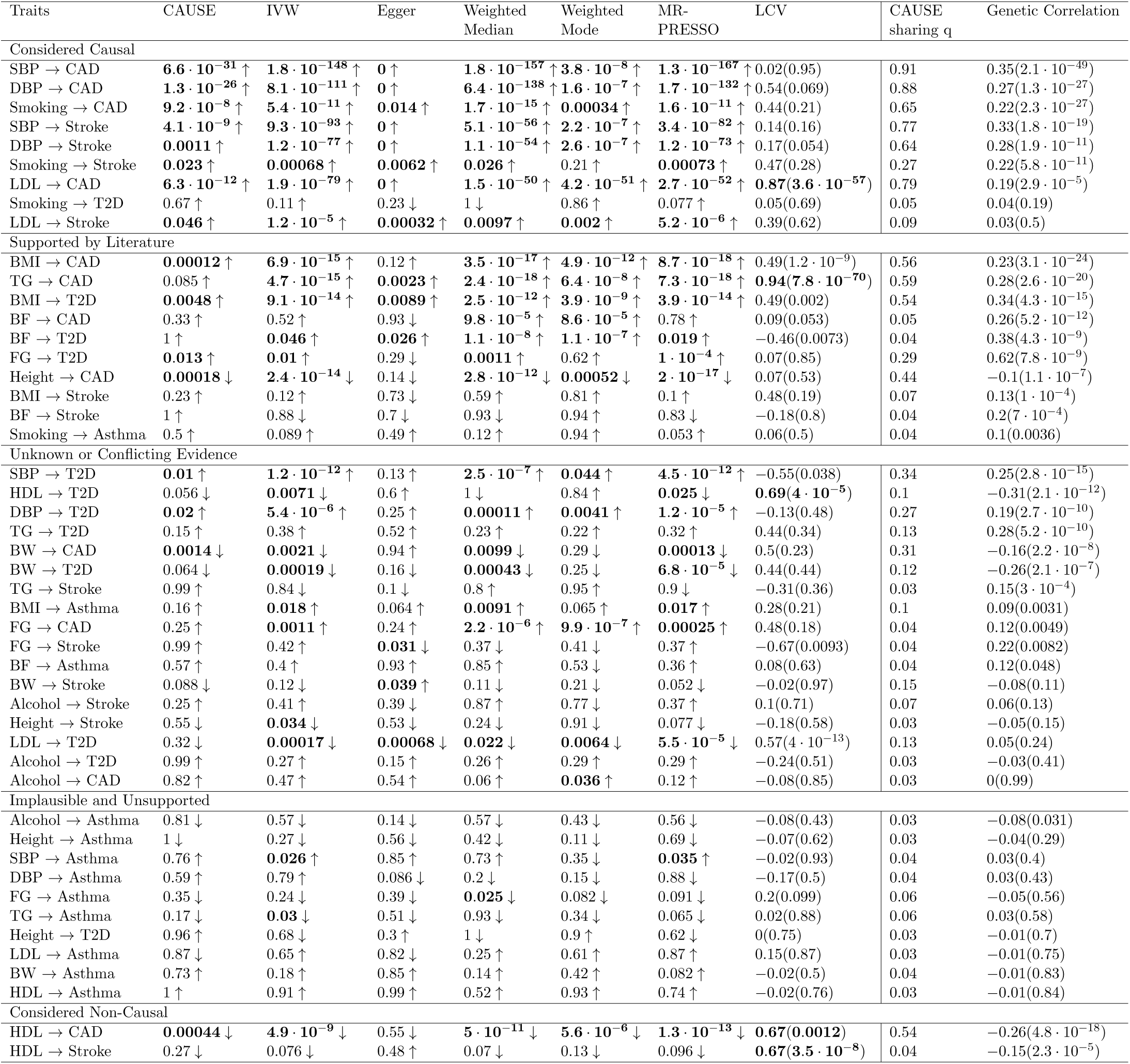
Summary of results for pairs of GWAS traits categorized by prior information about causality. Columns 2-7 give the *p*-value for each MR method. Values are bold if *p* < 0.05. Arrows indicate the sign of the corresponding effect estimate. The LCV column gives the estimated GCP from LCV with *p*-value testing that GCP= 0 in parenthesis. Values are bold if estimated GCP> 0.6. The “Cause *q*” column gives the posterior median of *q* in the CAUSE sharing model and the GC column gives the genetic correlation estimated by LD score regression with *p*-value testing that genetic correlation is zero in parenthesis. In each section, pairs are ordered by increasing genetic correlation *p*-value.

**Figure 6:**
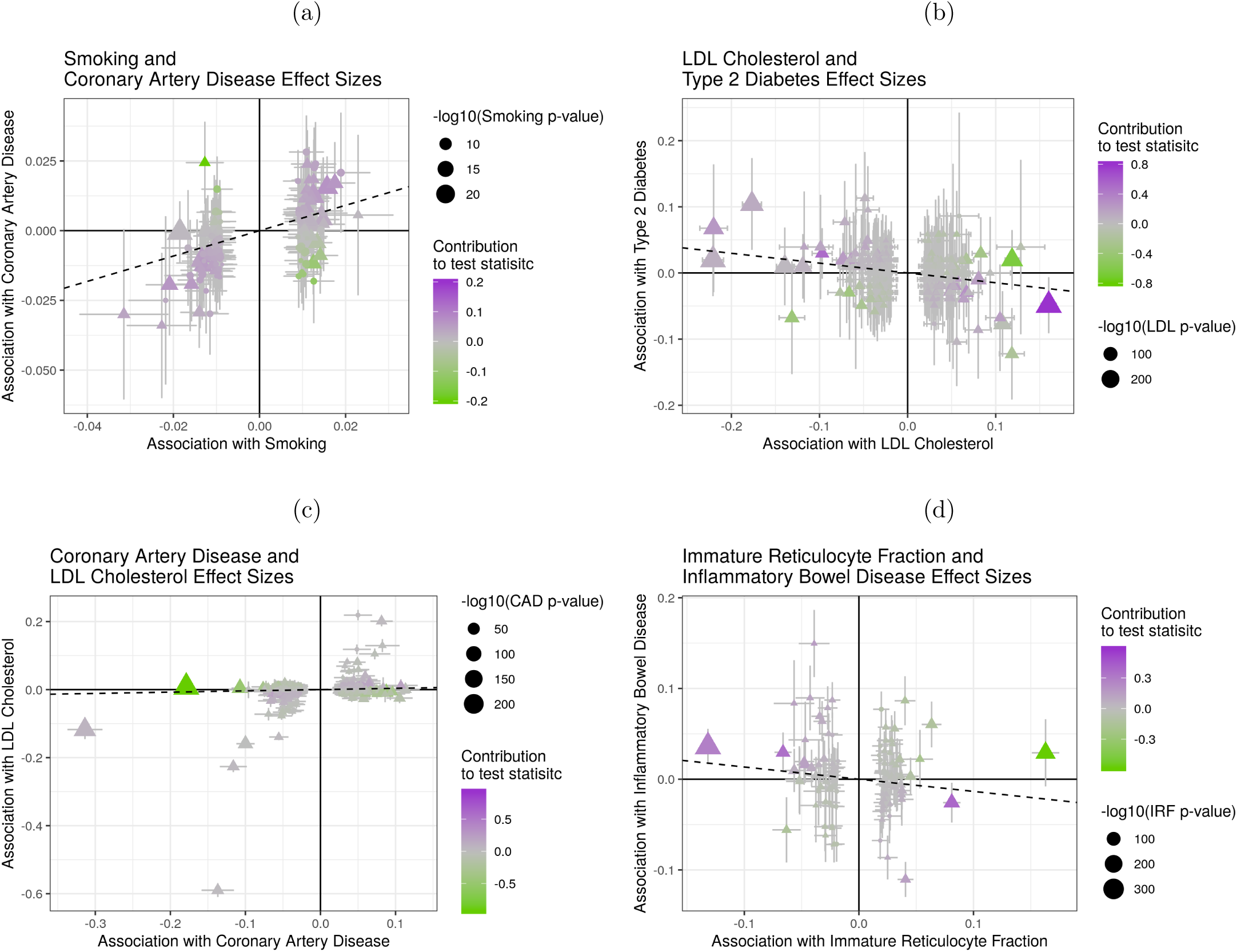
Effect size estimates and variant level contribution to CAUSE test statistics for four trait pairs. Effect estimates for trait *M* (horizontal axis) are plotted against estimates for trait *Y* (vertical axis). Error bars have length 1.96 times the standard error of the estimate. Triangles indicate variants reaching genome-wide significance for trait *M* (*p* < 5 · 10^−8^). Variants with trait *M p*-value < 5 · 10^−6^ are shown. Dotted lines show the IVW estimate obtained using only genome-wide significant variants. (a) Smoking (*M*) and CAD (*Y*). All methods detect evidence of a causal effect. (b) LDL (*M*) and T2D (*Y*). CAUSE does not detect a causal effect. Under the CAUSE model, these data can be explained by a shared factor associated with 13% of LDL effect variants. (c) CAD (*M*) and LDL cholesterol (*Y*). Some strong CAD variants have correlated effects on LDL, likely acting on CAD through effects on LDL. These lead to false positives for Egger regression, the weighted mode, and MR-PRESSO (*p* = 4.8 · 10^−5^, 0.0068 and 0.0013 respectively). CAUSE avoids a false positive (*p* = 0.11) by considering evidence from the many strong CAD variants that are not associated with LDL. IVW regression and the weighted median achieve *p*-values of 0.065 and 0.22. (d) IRF (*M*) and IBD (*Y*). MR-PRESSO, the weighted median, and modal estimators obtain a positive result by down-weighting or removing variants supplying conflicting evidence.

To assess false positives, we consider traits in the non-causal and implausible categories, and those in the unknown category that have no significant genetic correlation. CAUSE identifies fewer of these pairs as causal than any other method, suggesting that it is avoiding more false positives. All methods except Egger regression detect an effect of HDL cholesterol on CAD risk, which is likely to be an error[21]. This example represents a limitation of CAUSE, as it is possible to obtain false positives when a high proportion of variants are shared. In this case, CAUSE estimates that 54% of variants act through a shared factor.

LDL cholesterol and T2D is an example of a pair of traits identified by other methods but not by CAUSE (Figure 6b). The true relationship between this pair of traits is unknown. Levels of LDL cholesterol are often elevated in individuals with T2D, an effect that is sometimes considered a consequence of insulin insensitivity [22] (see Supplementary Note Section SN7). The effects estimated by IVW regression and other methods are negative, opposite of what might be expected from observational associations, but consistent with previous findings of Fall et al. [23]. CAUSE does not reject the sharing model (*p* = 0.32) and estimates that 13% of LDL variants act through a shared factor.

We expect that most effects detected in the opposite direction, of diseases on potential mediators are false positives. Testing opposite direction effects therefore gives us another opportunity to evaluate the robustness of methods to false positives. Additionally, as discussed previously, a reciprocal result for CAUSE can arise from both causal and non-causal scenarios, making reverse direction analyses important to consider. At a *p*-value threshold of 0.05, CAUSE identifies five reverse pairs. These are effects of CAD and stroke on diastolic and systolic blood pressure and of T2D on fasting glucose. IVW, Egger regression, the weighted median, the weighted mode, and MR-PRESSO identify 15, 6, 11, 7, and 17 reverse effects respectively (Table S6, Figure S4). The case of CAD and LDL cholesterol provides an example of how CAUSE is able to avoid false positives from reverse effects (Figure 6c, full results in Table S6).

All of the mediators in this analysis have high powered GWAS with many genome-wide significant variants. Such studies are not always available and simulations suggest that CAUSE may have a power advantage over other methods if the trait *M* GWAS is under-powered. In order to explore the effect of GWAS power, we compared results using an earlier GWAS for blood pressure that has a much smaller sample size [24]. With these data, IVW regression and Egger regression are unable to detect the effect of blood pressure on CAD. CAUSE is able to detect the effect by incorporating information from less significant variants (Supplementary Note Section SN9). This suggests that CAUSE may be able to boost power to detect causal effects of mediators with under-powered GWAS.

### 2.5 Links between immune-mediated disease and blood cell counts

There are a growing number of GWAS for intermediate or molecular traits. Relationships between these traits and clinical outcomes are appealing targets for MR studies because they may be accessible as drug targets and may suggest molecular and physiological pathways contributing to disease. We use CAUSE and other MR methods to search for evidence of causal relationships between 13 measures of blood cell composition and five immune-mediated diseases: allergic disease, asthma, rhumatoid arthritis, inflammatory bowel disease, and systemic lupus erythematosus (Table S2). Results are summarized in Figure 7 and Table S4 (full results in Tables S7 and S8).

**Figure 7:**
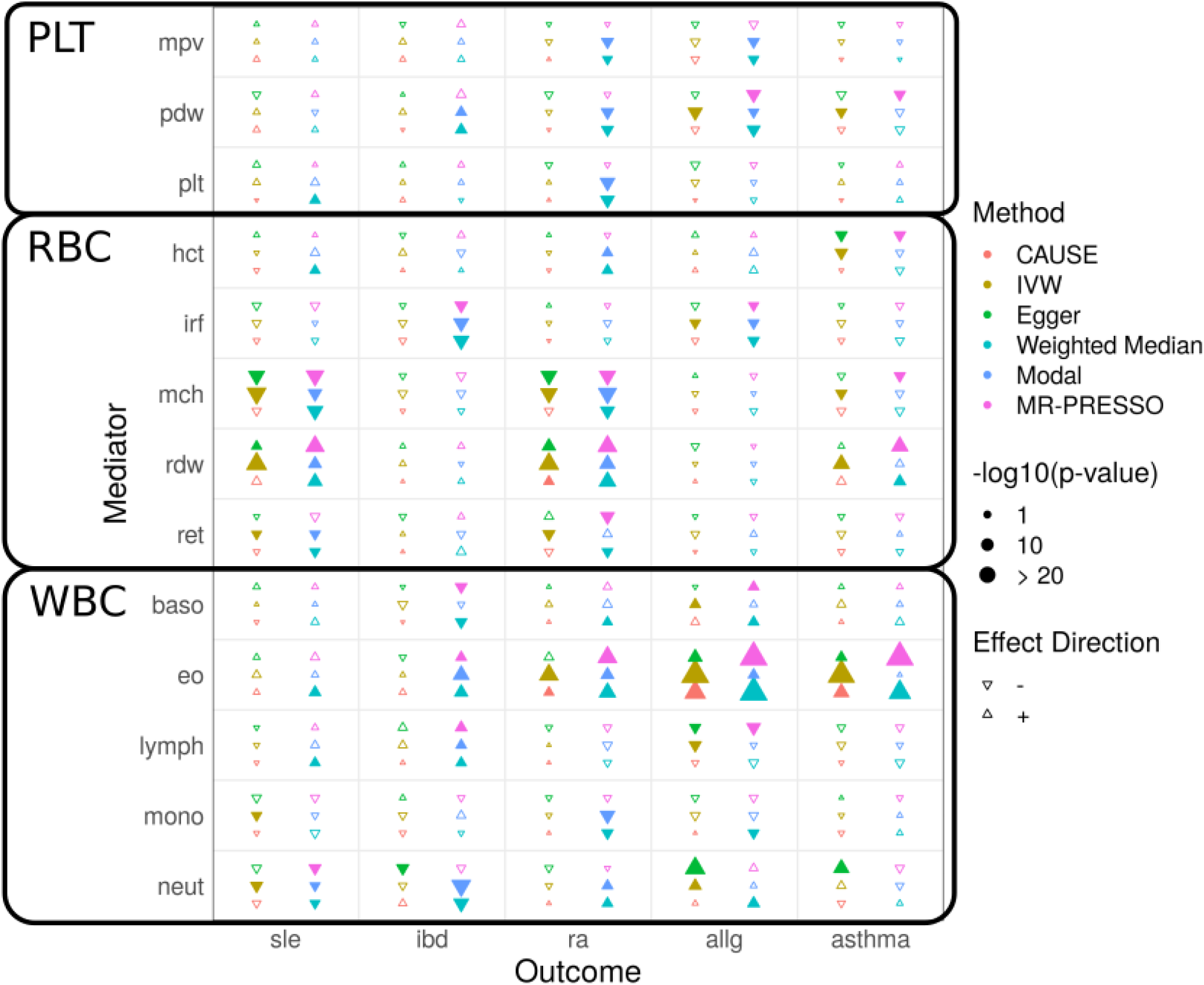
Tests for causal effects of blood cell composition on immune mediated traits. Each cell summarizes the the results of six methods for a pair of traits. Filled symbols indicate *p*-value < 0.05. Blood cell traits are grouped into platelet traits (PLT), red blood cell traits (RBC), and white blood cell traits (WBC).

The blood cell composition traits can be divided into red blood cell traits, white blood cell traits, and platelet traits. There is less literature information about which relationships are expected to be causal than in the previous application. However, we expect that relationships between immune-mediated disorders and immune related white blood cell traits are more plausible than relationships with red blood cell or platelet traits.

CAUSE identifies only 4 pairs of traits as consistent with a causal effect at a *p*-value threshold of 0.05. The two most significant effects are positive effects of eosinophil count on asthma and allergy risk (*p* = 1.9 · 10^−5^ and 4.0 · 10^−11^ respectively). Eosinophils have proven to be promising drug targets for treating asthma and allergic disease [25]. All other methods, including Egger regression and the weighted mode obtain many more positive results than CAUSE, including many causal relationships of red blood cell and platelet traits which are likely false positives. Figure 6d shows a pair of traits, immature reticulocyte fraction (IRF, a red blood cell trait) and inflammatory bowel disease (IBD), that are only detected by methods designed to be robust to pleiotropy, the weighted median (*p* = 1.2 · 10^−5^), the weighted mode (*p* = 2.6 · 10^−5^), and MR-PRESSO (*p* = 0.0015). There is no visible correlation in variant effect sizes and CAUSE as well as the simpler IVW regression obtain non-significant results (*p* = 0.32 and 0.16 respectively). However, by removing or down-weighting the contribution of some variants, these methods can obtain false positives by discounting conflicting evidence.

## 3 Discussion

We have introduced CAUSE, a new approach to MR analysis that accounts for uncorrelated and correlated pleiotropy. CAUSE has several advantages over existing methods, most notably reduction of false positives. Previous authors have identified uncorrelated horizontal pleiotropy as a pervasive phenomenon that may adversely impact MR analyses [7]. Our analyses demonstrate that correlated horizontal pleiotropy is also common and may explain a substantial fraction of positive results obtained using other MR methods. Alternative MR methods perform best when information is available to guide removal of pleiotropic variants variants. For example, in analyzing HDL cholesterol and heart disease, Voight et al. avoid a false positive by discarding variants associated with other lipid and metabolic traits. CAUSE, on the other hand, is well suited for applications where it is difficult to predict shared factors, shared factors cannot be measured, or the researcher would like to screen a large number of potentially causal variables in an automated manner.

Caution must be used when interpreting results of CAUSE, as well as any other MR method. CAUSE tests that the GWAS summary statistics for *M* and *Y* are consistent with a model in which every *M* effect variant has a correlated effect on *Y*. This pattern occurs when *M* has a causal effect on *Y* but can also occur in other circumstances. Notably, if most of the heritable variation of two traits is mediated by the same unobserved process, we expect to observe this pattern in both trait directions. With this in mind, it is good practice to test trait pairs in both directions [13]. We also note that while CAUSE is more robust than alternatives, it still has a high false positive rate when the shared factor has a strong effect and accounts for a large proportion of the trait *M* variants. Thus, a low *p*-value from CAUSE (as with other MR methods) should not be regarded as proof of a causal effect. Instead, it is an indicator that the summary statistics for the two traits are consistent with a causal effect.

CAUSE has several limitations that provide interesting future research directions. First, it is not currently possible to account for known shared factors when they are measured. Second, CAUSE models only a single unobserved shared factor, so it may not fully account for shared genetic components between two traits of interest. This problem is partially alleviated by the flexibility of the empirical effect size prior distribution (see Methods). Finally, CAUSE simply prunes variants for LD rather than explicitly modeling variant correlation. This helps ensure that the problem is computationally tractable, however, using only one variant per LD block may lead to loss of information.

## 4 Methods

### 4.1 CAUSE model for GWAS Summary Statistics

We use effect estimates and standard errors measured in GWAS of *M* and *Y* (summary statistics) to evaluate evidence of a causal effect of *M* on *Y*. Let 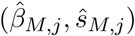 and 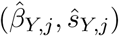 be effect estimates and standard errors at variant *G*_*j*_ (*j* = 1, …, *p*) for traits *M* and *Y* respectively. Let *β*_*M,j*_ and *β*_*Y,j*_ be the true marginal associations of *G*_*j*_ with *M* and *Y*. We model effect estimates as normally distributed given the true effects, allowing for global correlation that can result from overlapping GWAS samples. We model

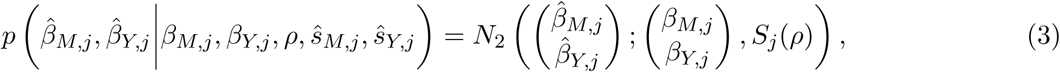

where where 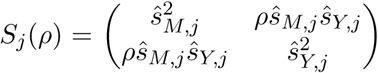 and *N*_2_(*x*; *µ*, Σ) is the bivariate normal density with mean *µ* and variance Σ evaluated at *x*. This model implicitly assumes that ŝ_*M,j*_ and ŝ_*Y,j*_ are measured without error. The correlation term, *ρ*, which accounts for sample overlap, is estimated empirically (Supplementary Note Section SN1).

In the CAUSE model, a proportion, *q*, of variants exhibit correlated pleiotropy, modeled as an effect on a shared factor, *U* (Figure 3b). The remaining proportion, 1 − *q*, are independent of *U* (Figure 3a). All variants may have pleiotropic effects, *θ*_*j*_ on *Y* that are uncorrelated with their effects on *M*. Let *Z*_*j*_ be an indicator that variant *j* effects *U*. Then,

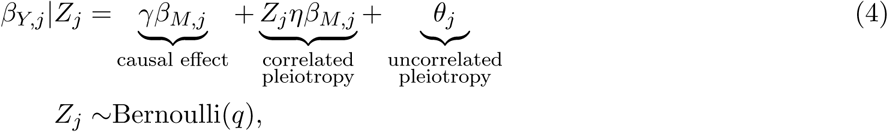

where *η* is the effect of *U* on *Y* and *U* is scaled so that the effect of *U* on *M* is 1. Note that if there are no uncorrelated or correlated pleiotropic effects, then Equation (4) reduces to *β*_*Y,j*_ = *γβ*_*M,j*_, which is the relationship assumed by simple MR approaches such as IVW regression. We substitute the right side of Equation (4) into Equation (3) and integrate out *Z*_*j*_ to obtain

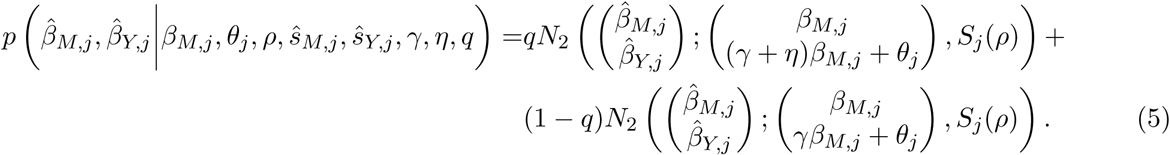

The parameters of interest in the CAUSE model are *γ, η*, and *q*. Rather than estimating individual variant effects *β*_*M,j*_ and *θ*_*j*_, we model their joint distribution empirically and integrate them out to obtain a marginal density for 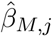 and 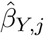. We model *β*_*M,j*_ and *θ*_*j*_ as draws from a mixture of bivariate normal distributions. This strategy is based on Adaptive Shrinkage (ASH) approach of Stephens [26] for modeling univariate distributions and provides a flexible unimodal distribution with mode at (0, 0). We model

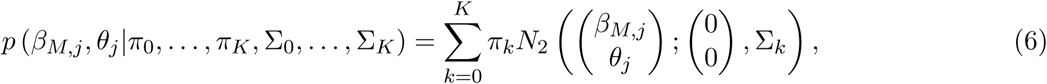

where Σ_0_, Σ_1_, …, Σ_*K*_ are pre-specified covariance matrices of the form 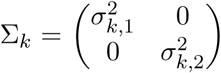, and *π*_0_, …, *π*_*K*_ are mixing proportions that sum to 1. The set of parameters Ω = {*π*_0_, …, *π*_*K*_, Σ_0_, …, Σ_*K*_} are estimated from the data along with *ρ* in a single pre-processing step (Supplementary Note Section SN1). Integrating *β*_*M,j*_ and *θ*_*j*_ out of Equation (5), we obtain

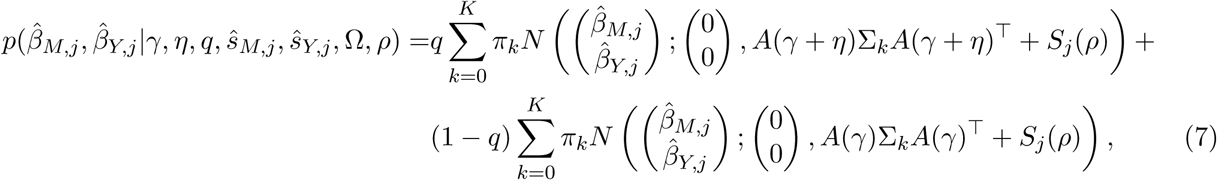

where 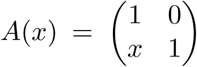. Treating variants as independent, we obtain a joint density for the entire set of summary statistics as a product over variants. We place independent prior distributions on *γ, η*, and *q*, described below, and estimate posterior distributions via adaptive grid approximation (Supplementary Note Section SN3). A flow-chart illustrating a CAUSE analysis including parameter estimation is shown in Figure S1. CAUSE is implemented in an open source R package and we also provide pipelines for running CAUSE on large scale applications.

### 4.2 Prior Distributions for *γ, η*, and *q*

We place normal prior distributions with mean zero on *γ* and *η*. We find in simulations that results are robust to different choices of prior variance for *γ* and *η*, as long the same value is used for both parameters (Supplementary Note Section SN2). Since the magnitude of a possible causal effect is difficult to know a priori due to differences in trait scaling and covariate adjustment across GWAS, we use the data to suggest a prior variance (Supplementary Note Section SN2).

By default, and in all analyses presented here, we use a Beta(1, 10) prior distribution for *q*. This distribution gives a prior probability of 0.001 that *q* > 0.5 and 0.056 that *q* > 0.25. The parameters of the Beta distribution can be adjusted by the user to reflect different prior beliefs about the size of *q*. If the prior on *q* places more weight on values close to zero, CAUSE will behave more similarly to standard MR, obtaining high false positive rates when the true value of *q* is large. With a more permissive prior on *q*, CAUSE is more robust to larger amounts of correlated pleiotropy but has reduced power. We set Beta(1, 10) as the default choice because it performed acceptably over a range of settings in simulations (Supplementary Note Section SN6.1).

### 4.3 Accounting for Linkage Disequilibrium

We treat variants as independent when we compute the joint density of summary statistics. In reality, variants are correlated due to linkage disequilibrium (LD). We define the *LD-transformed effects* 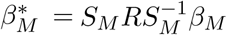 and 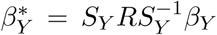, where *R* is the variant correlation matrix matrix and *S*_*M*_ and *S*_*Y*_ are diagonal matrices with diagonal elements (*s*_*M,j*_) and (*s*_*Y,j*_) [19]. We assume that *R* is the same in both GWAS. The pair of estimates 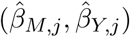 can be approximated as normally distributed with mean 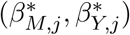 and variance *S*(*ρ*). If the relationship in Equation (4) holds for true effects, *β*_*M,j*_ and *β*_*Y,j*_, and either *M* effects are sparse relative to LD structure (most *M* effect variants are independent) or *q* is small, then the relationship between the LD-transformed effects is

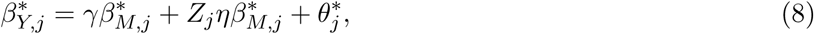

where 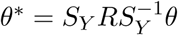 (Supplementary Note Section SN4). In this case, the mean relationship for summary statistics at a single locus is the same with or without LD and Equation (5) and (7) remain valid for variants in LD if *β*_*M,j*_ and *θ*_*j*_ are replaced with 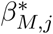 and 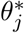.

Correlations among variants can affect the joint density of all summary statistics, which we compute as a product of the densities at each variant. To account for this, we use a subset of variants with low mutual LD (*r*^2^ < 0.1 by default), prioritizing variants with low trait *M p*-values to improve power.

If *M* effect variants are dense relative to LD structure and *q* is large, LD can induce a positive correlation between *Y* and *M* effect estimates for all variants even when there is no causal effect, leading to false positives (Supplementary Note Section SN4). In this case, CAUSE may have an inflated false positive rate.

### 4.4 Model Comparison Using ELPD

To determine whether GWAS summary statistics are consistent with a causal effect of *M* on *Y* we compare a model in which the causal effect is fixed at zero (the *sharing model*) to a model that allows a non-zero causal effect (the *causal model*). To compare the fit of these models, we estimate the difference in the *expected log pointwise posterior density* (ΔELPD) [18]. The ELPD measures how well the posterior distributions estimated under a given model are expected to predict a hypothetical new set of summary statistics obtained from GWAS of *M* and *Y* in different samples.

Let Θ be the set of parameters (*γ, η, q*). Let *p*_*C*_ (Θ|Data) and *p*_*S*_ (Θ|Data) be the posterior density of Θ given the observed summary statistics under the causal model and sharing model respectively. Let 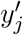 denote a new observation of 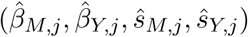. The ELPD for model *m* ∈ {*C, S*} (for causal and sharing) is

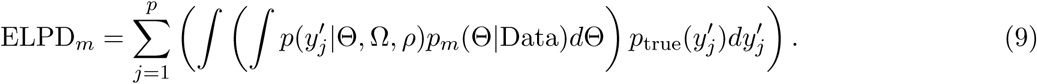

where 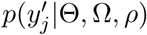 is given in Equation (7) and *p*_true_ is the probability density under the true data generating mechanism. If ΔELPD = ELPD_*C*_ − ELPD_*S*_ is positive, then the posteriors from the causal model predict the data better so the causal model is a better fit. If ΔELPD≤ 0, then the sharing model fits at least as well, indicating that the data are not consistent with a causal effect.

We estimate ΔELPD and a standard error of the estimator using the Pareto-smoothed importance sampling method described by Vehtari, Gelman, and Gabry [18] and implemented in the R package loo. We then compute a *z*-score, 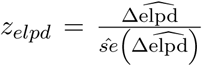, that is larger when posteriors estimated under the causal model fit the data better than the posteriors estimated under the sharing model. We compute a one-sided *p*-value by comparing the *z*-score to a standard normal distribution. The *p*-value estimates the probability of obtaining a *z*-score larger than the one observed if the true value of ΔELPD were less than or equal to zero.

### 4.5 Generating Simulated Summary Statistics

To create data with a realistic correlation structure, we estimate LD for 19,490 HapMap variants on chromosome 19 in the CEU 1,000 Genomes population using LDShrink ([27]; https://github.com/stephenslab/ldshrink) and replicate this pattern 30 times to create a genome sized data set of *p* = 584, 700 variants. We set the heritability of each trait, 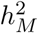 and 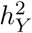, the sample size for each GWAS, *N*_*M*_ and *N*_*Y*_, and the expected number of effect variants *m*_*M*_ and *m*_*θ*_. Additionally, we set *γ, η*, and *q* and generate effect estimates from the CAUSE model in Equation (4). We note that *m*_*θ*_ is the expected number of variants with *θ*_*j*_ ≠ 0, rather than the expected number of variants with non-zero effect on *Y*.

We simulate an effect estimate and standard error for each variant using the following procedure. First, standardized effects 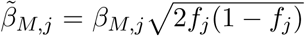 and 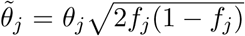 are drawn from a mixture distribution:

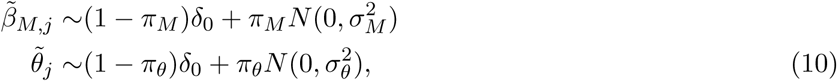

where *f*_*j*_ is the allele frequency of variant *j* in the 1,000 Genomes CEU population. Variances 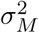 and 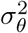 chosen to give the desired expected heritability as 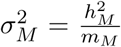 and 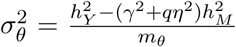. Mixing parameters are *π*_*M*_ = *m*_*M*_ */p* and *π*_*θ*_ = *m*_*θ*_*/p*. Note that 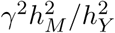 is the proportion of trait *Y* heritability mediated by the causal effect and 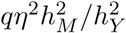 is the proportion of trait *Y* heritability mediated by *U*.

Second, standardized effects are converted to non-standardized effects and standard errors are computed as as 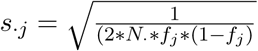, where may be *M* or *Y*. Third, *Z*_*j*_ are drawn from a Bernoulli(*q*) distribution and true effects *β*_*γ,j*_ are computed using Equation (4). Finally, effect estimates are simulated from true effects as

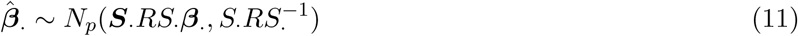

where *S*_·_ is the diagonal matrix of standard errors and *R* is the variant correlation matrix [19]. Simulations can be replicated using the online tutorial (https://jean997.github.io/cause/simulations.html).

### 4.6 Existing MR Methods

We compare the performance of CAUSE in simulated data with six other MR methods and LCV [14]. These are implemented as follows.

- Random effect IVW regression, random effect Egger regression, the weighted median, and weighted mode methods are implemented in the `MendelianRandomization` R package. IVW regression is run using `model = “random”` and `weights = “delta”` options. The weighted mode estimator is computed using the `phi = 1` and `stderror=“delta”` options. All other methods are run with default options.
- MR-PRESSO is performed using the `MRPRESSO` R package ([7], https://github.com/rondolab/MR-PRESSO) with outlier and distortion tests.
- GSMR is performed using the `gsmr` R package ([10], http://cnsgenomics.com/software/gsmr/) with the Heidi outlier test, default threshold 0.01, and minimum number of instruments lowered to 1.
- LCV is performed using R scripts available from the author ([14], https://github.com/lukejoconnor/LCV) using default parameters.

### 4.7 URLs and Software Availability

All software and analysis code is publicly available. The CAUSE method is implemented in an R package available through GitHub.

- Website: https://jean997.github.io/cause/
- Includes pipelines and instructions for replicating all results presented in this paper.
- CAUSE Software (R package): https://github.com/jean997/cause
- Simulations Software (R package): https://github.com/jean997/causeSims

## 4.8 Acknowledgements

This work was supported by National Institutes of Health (NIH) grants MH110531 (to XH) and HG002585 (to MS), and a Research Grant from the March of Dimes (XH).

## 4.9 Data Availability

All of the data analyzed are publicly available with the exception of blood pressure summary statistics from [24]. These are available through dbGaP Accession phs000585.v2.p1. Download links for all other data sets as well as instructions for formatting and processing data and reproducing CAUSE analysis results can be found on the website https://jean997.github.io/cause/.

## Supplementary Figures

**Figure S1:**
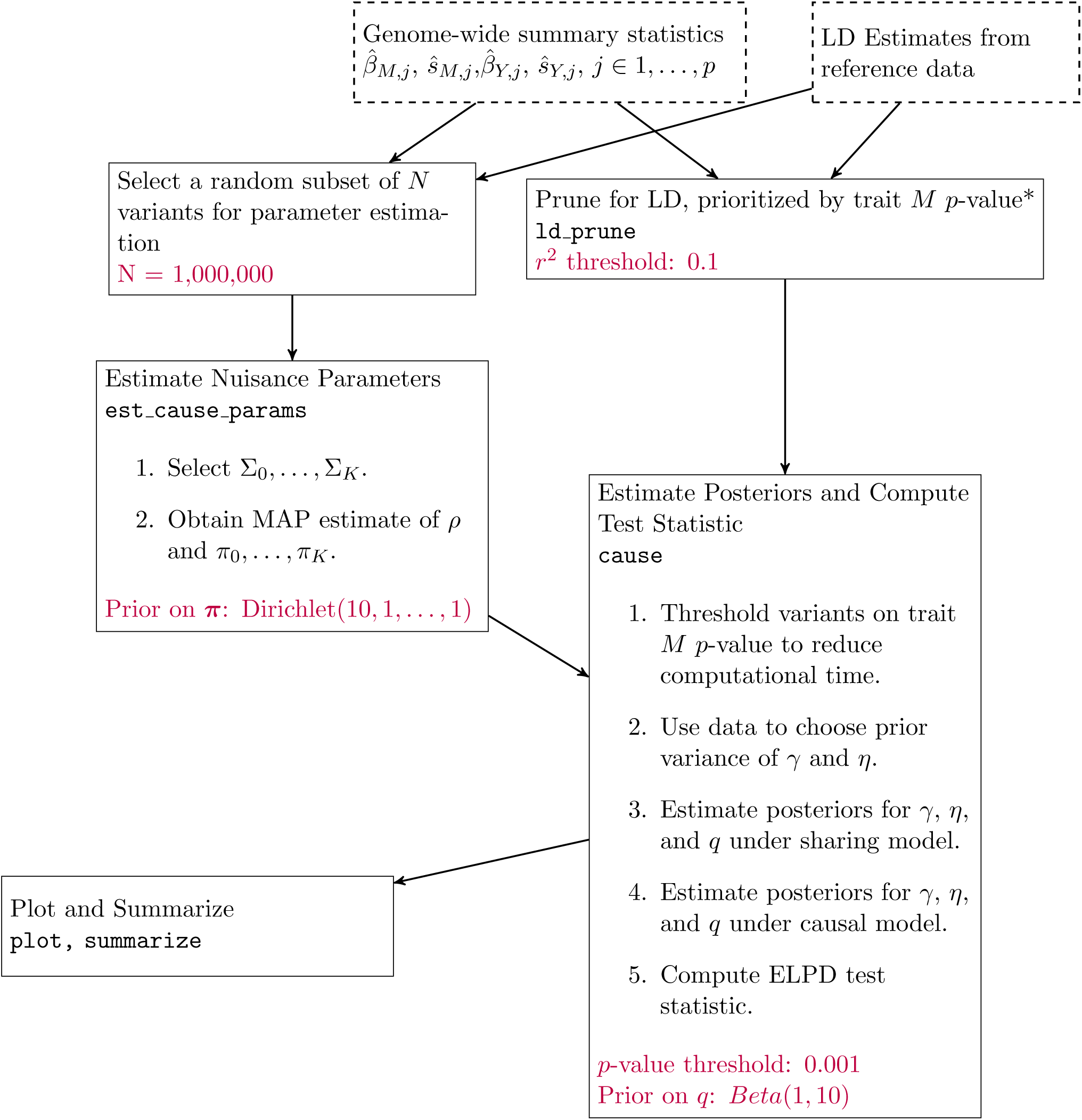
Workflow of a CAUSE analysis. Dashed boxes represent input data. Each solid box is an analysis step completed by the given function in the cause R package. LD pruning (*) can be parallelized over chromosomes. Purple text indicates user provided parameters and their default values. All analyses presented are run with default parameters.

**Figure S2:**
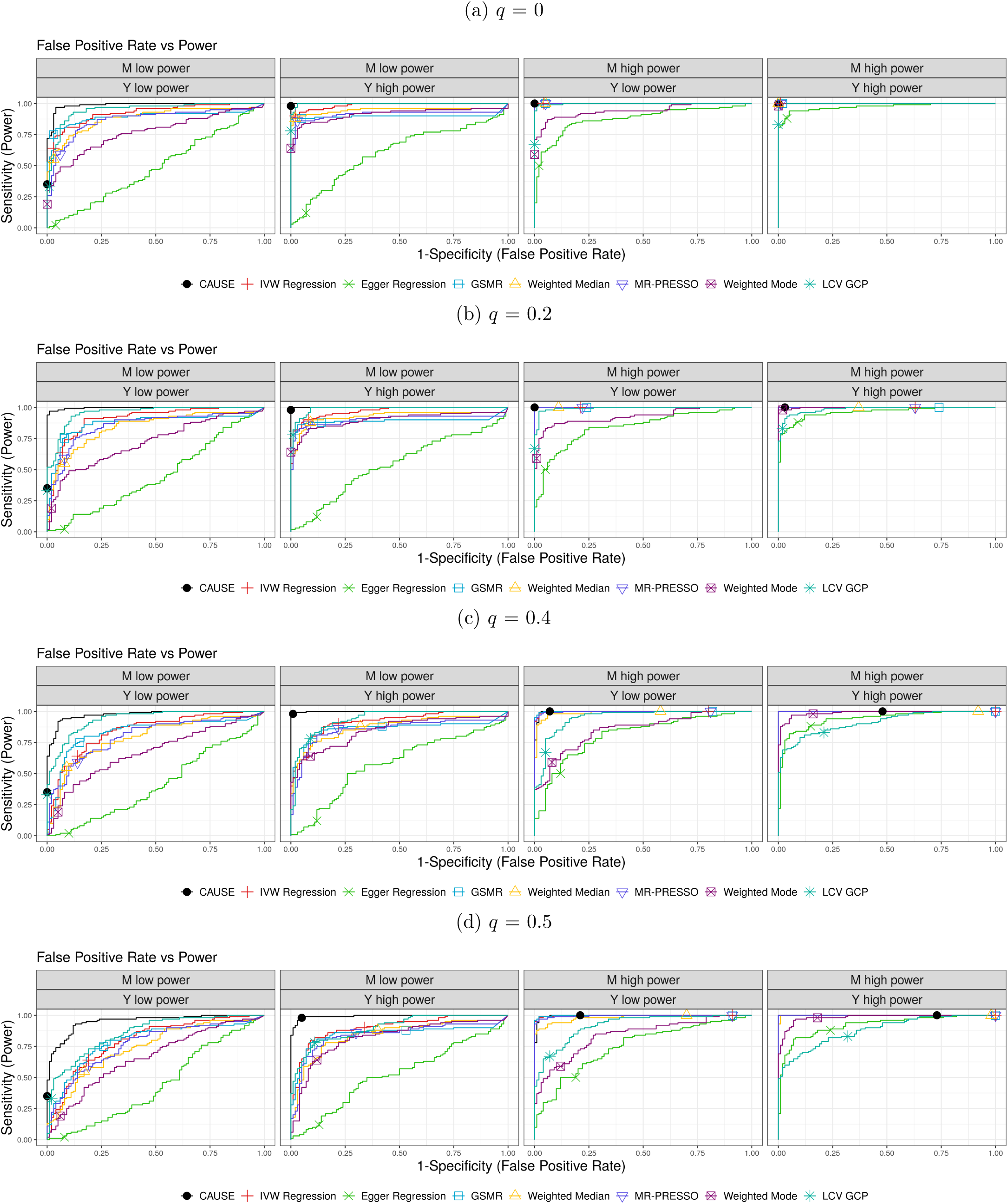
Comparison of false positive-power trade-off. We compare the power when 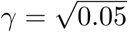 and there is no shared factor to the false positive rate when there is no causal effect, but a proportion of variants between 0 and 50% act through a shared factor with effect 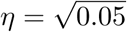 on *Y*. There are 100 simulations each in the causal and non-causal scenarios. Curves are created by varying the significance threshold. Points indicate the power and false positive rate achieved at a threshold of *p* ≤ 0.05.

**Figure S3:**
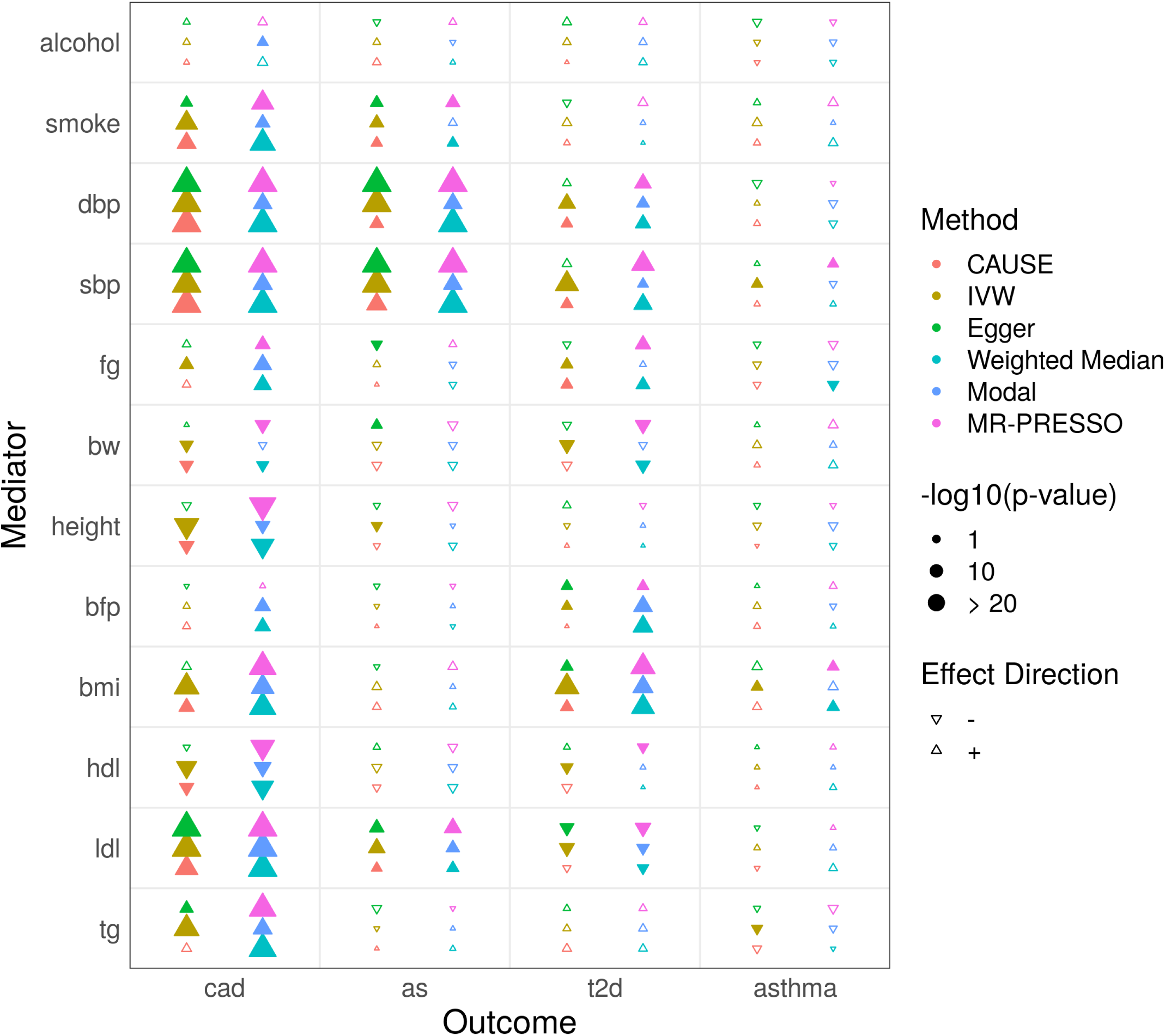
Tests for casual effects of mediators on diseases. Each cell summarizes the the results of six methods for a pair of traits. In the left column of the cell methods from bottom to top are CAUSE, IVW regression, and Egger regression. In the right column methods from bottom to top are weighted median, weighted mode, and MR-PRESSO. Filled symbols indicate a nominally significant *p*-value < 0.05.

**Figure S4:**
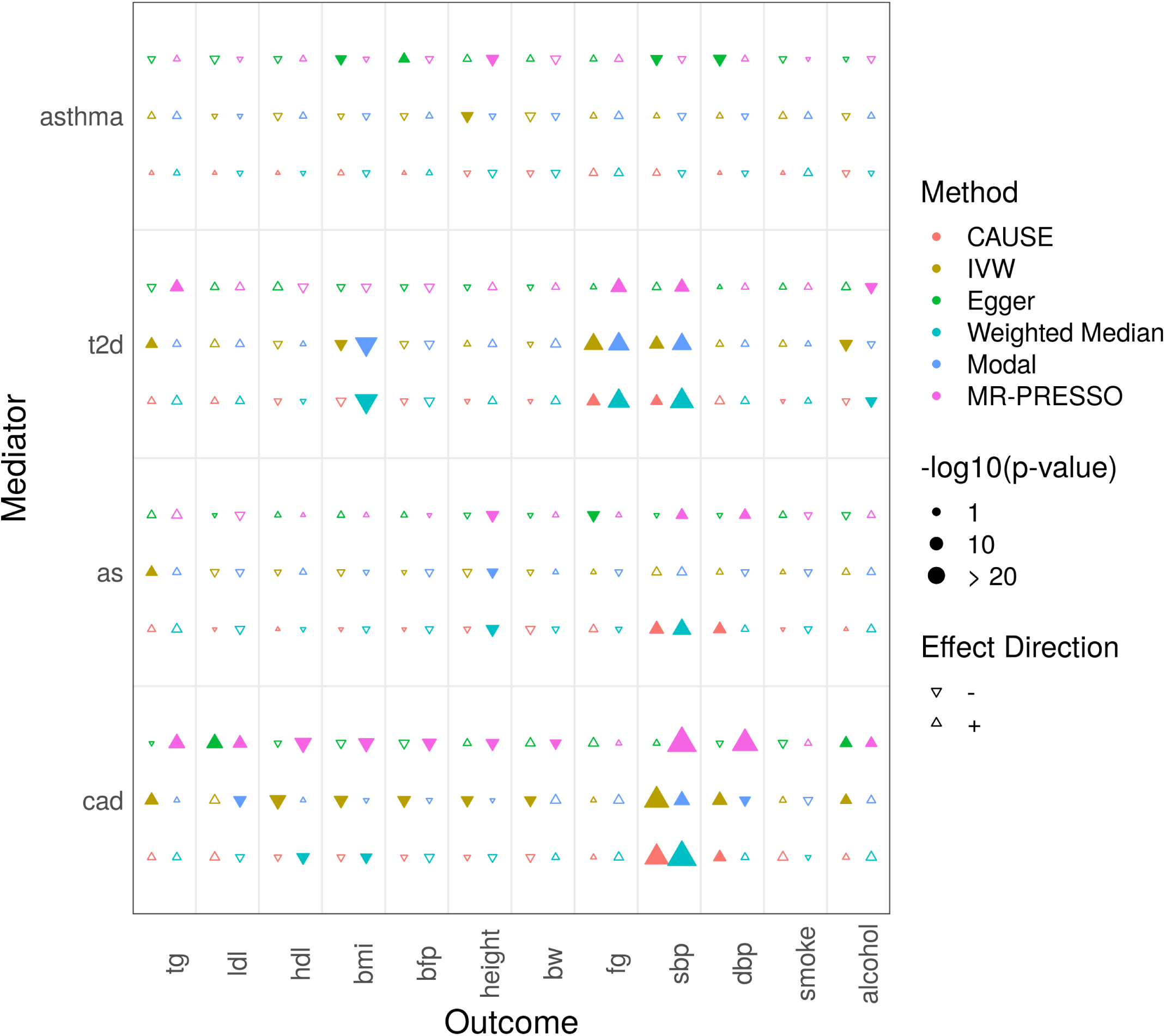
Tests for casual effects of disease outcomes on mediators. Each cell summarizes the the results of six methods for a pair of traits. In the left column of the cell methods from bottom to top are CAUSE, IVW regression, and Egger regression. In the right column methods from bottom to top are weighted median, weighted mode, and MR-PRESSO. Filled symbols indicate a nominally significant *p*-value < 0.05.

**Figure S5:**
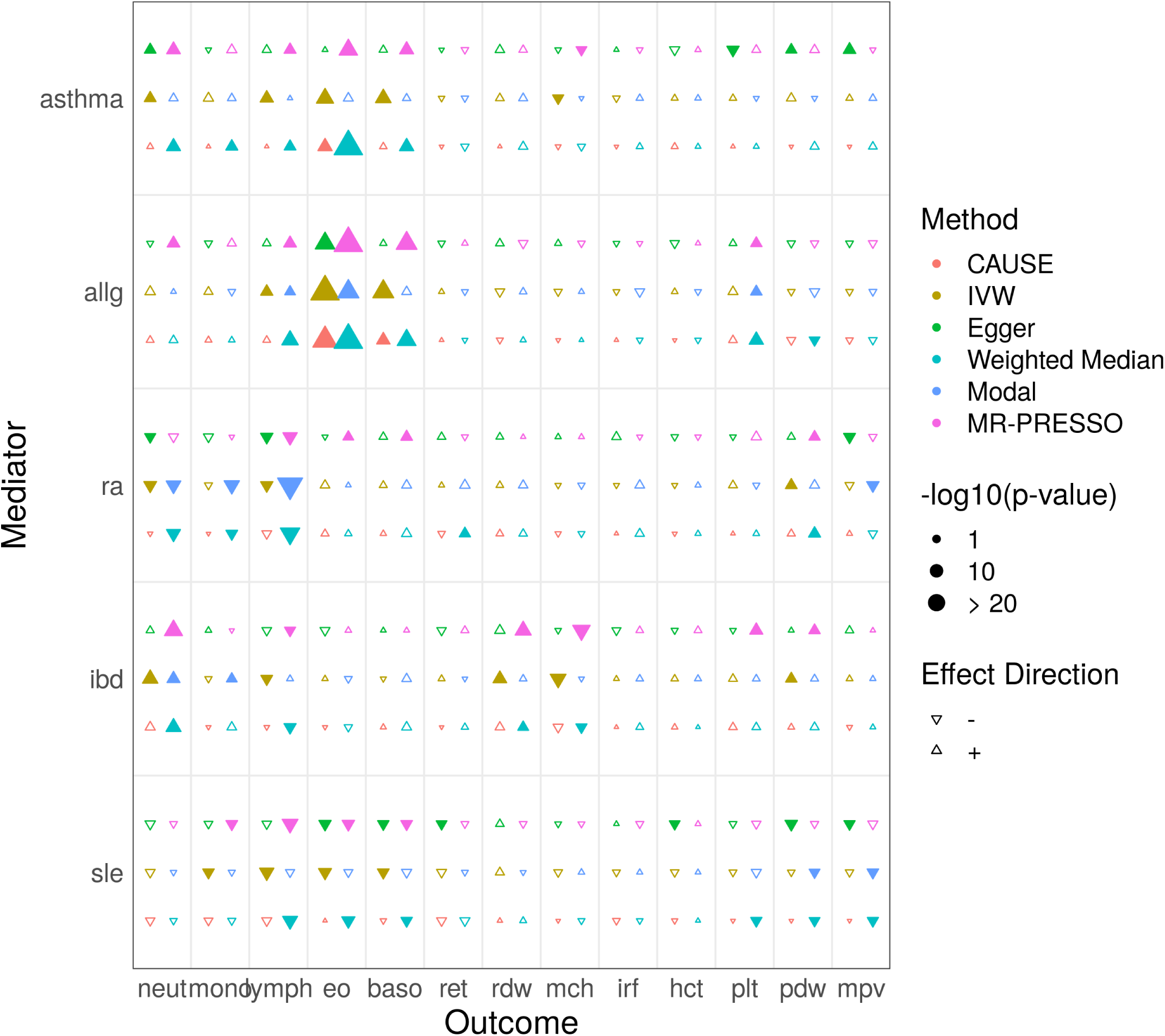
Tests for causal effects of autoimmune traits on blood cell composition. Each cell summarizes the the results of six methods for a pair of traits. In the left column of the cell methods from bottom to top are CAUSE, IVW regression, and Egger regression. In the right column methods from bottom to top are weighted median, weighted mode, and MR-PRESSO. Filled symbols indicate a nominally significant *p*-value < 0.05.

## Supplementary Tables

**Table S1:** Genome wide association studies for common diseases and risk factors

| Abbreviation | Trait | Sample Size | Cases | Controls | PMID | First Author (Year) |
| --- | --- | --- | --- | --- | --- | --- |
| tg | triglycerides | 188577 |  |  | 24097068 | Willer (2013) |
| ldl | ldl | 188577 |  |  | 24097068 | Willer (2013) |
| hdl | hdl | 188577 |  |  | 24097068 | Willer (2013) |
| height | height | 253288 |  |  | 25282103 | Wood (2014) |
| bmi | body mass index | 322154 |  |  | 25673413 | Locke (2015) |
| bf | body fat percentage | 100716 |  |  | 26833246 | Lu (2016) |
| bw | birth weight | 153781 |  |  | 27680694 | Horikoshi (2016) |
| dbp | diastolic blood pressure | 757601 |  |  | 30224653 | Evangelou (2018) |
| sbp | systolic blood pressure | 757601 |  |  | 30224653 | Evangelou (2018) |
| fg | fasting glucose | 46186 |  |  | 20081858 | Dupuis (2010) |
| smoke | ever regular smoker | 1232091 |  |  | 30643251 | Liu (2019) |
| alcohol | drinks per week | 941280 |  |  | 30643251 | Liu (2019) |
| cad | coronary artery disease | 547261 | 122733 | 424528 | 29212778 | van der Harst (2017) |
| stroke | any stroke | 446696 | 40585 | 406111 | 29531354 | Malik (2018) |
| t2d | type 2 diabetes | 69033 | 12171 | 56862 | 22885922 | Morris (2012) |
| asthma | asthma | 142486 | 23948 | 118538 | 29273806 | Demenais (2018) |

**Table S2:**
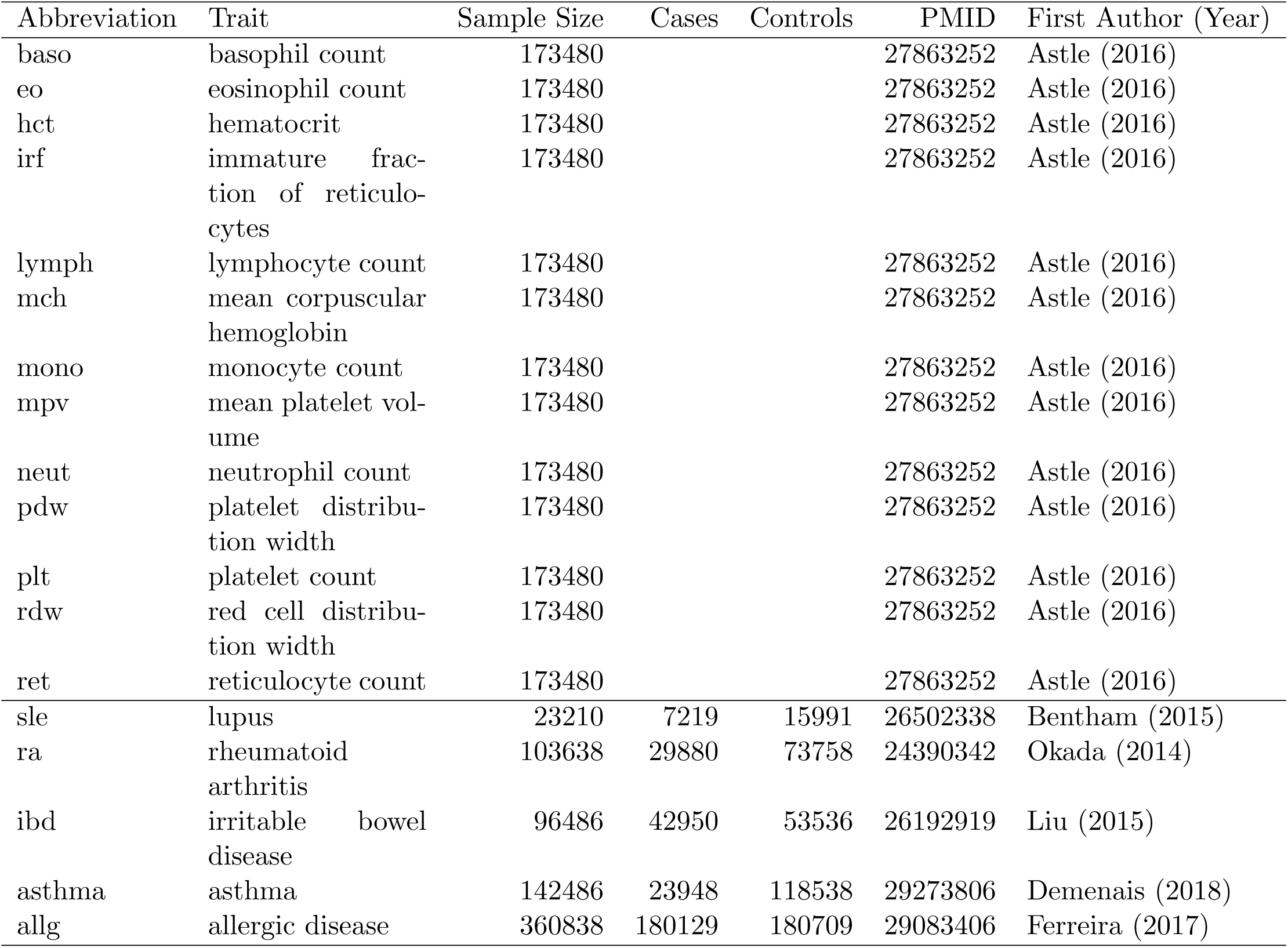
Genome wide association studies for blood cell traits and immune-mediated disesase.

**Table S3:**
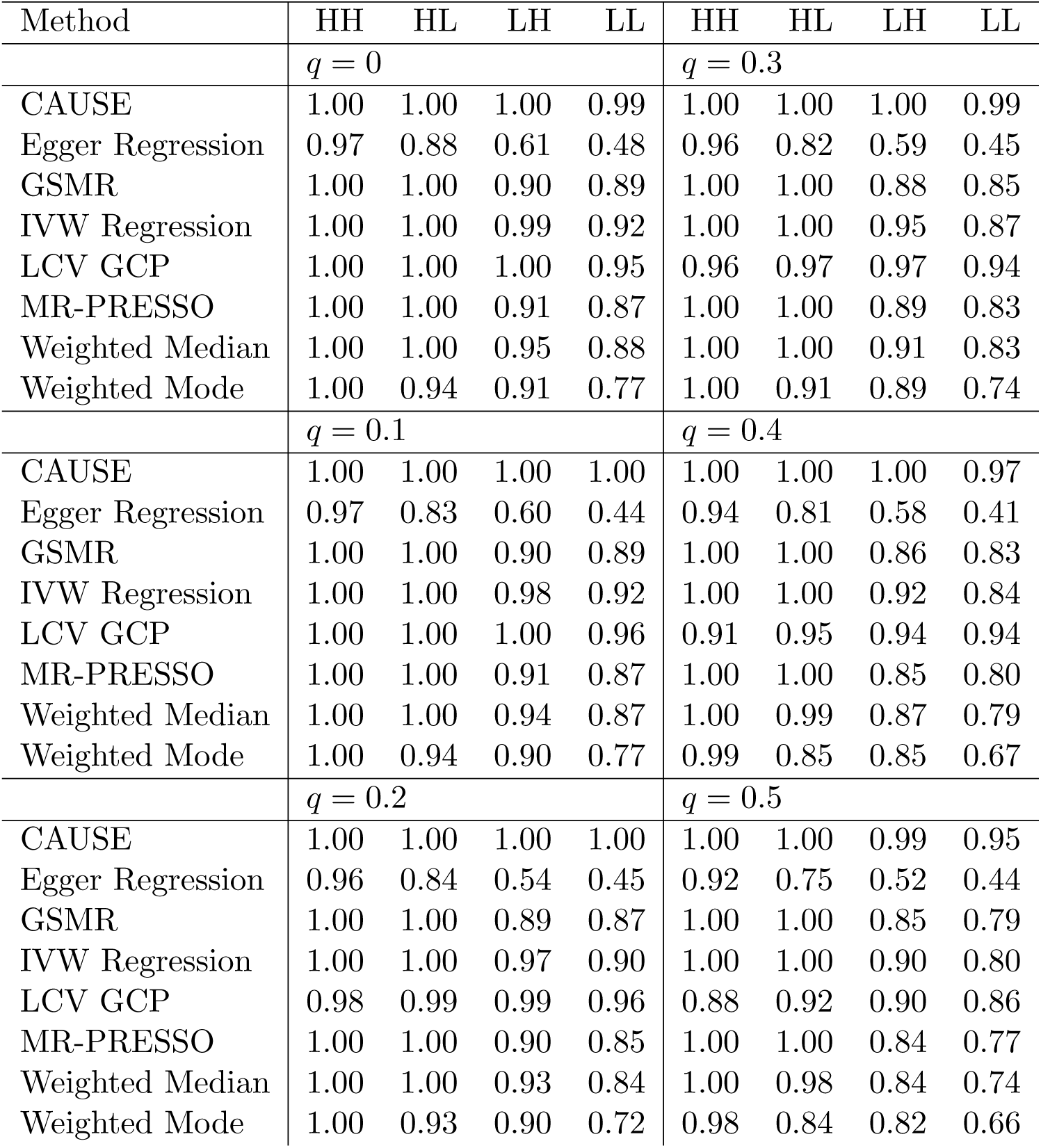
Area under curve values for CAUSE and other methods for ROC curves shown in Figures 4c and Supplementary Figure S2. Column labels (HH, HL, LH, and LL) indicate the power of trait *M* and *Y* GWAS (H = high power, L = low power) with the first letter referring to trait *M* and the second letter referring to trait *Y*.

**Table S4:**
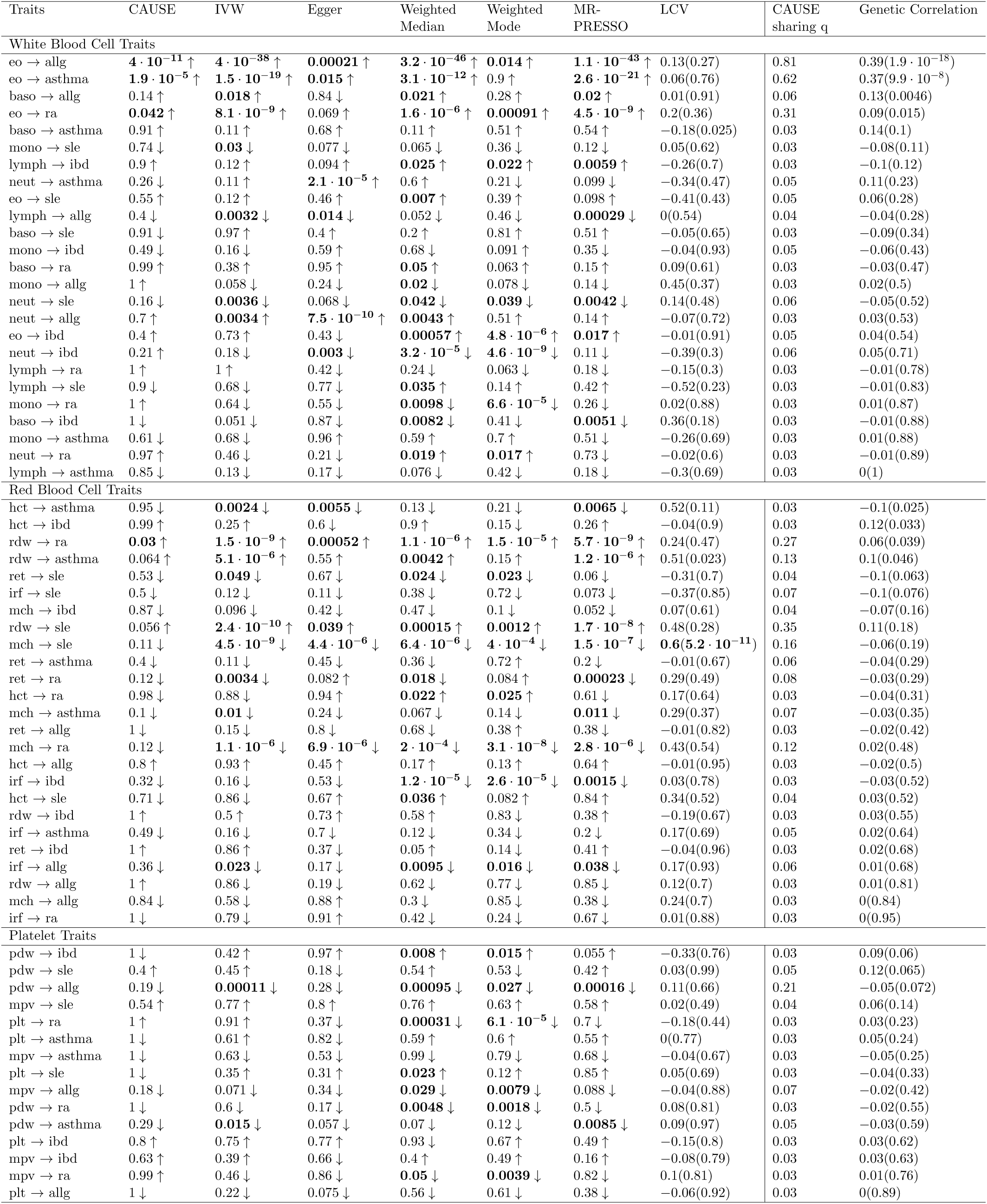
Summary of results for blood cell traits and immune-mediated diseases grouped by blood cell category. Columns 2-7 give the *p*-value for each MR method. Values are bold if *p* < 0.05. Arrows indicate the sign of the corresponding effect estimate. The LCV column gives the estimated GCP from LCV with *p*-value testing that GCP= 0 in parenthesis. Values are bold if estimated GCP> 0.6. The “Cause *q*” column gives the posterior median of *q* in the CAUSE sharing model and the GC column gives the genetic correlation estimated by LD score regression with *p*-value testing that genetic correlation is zero in parenthesis. In each section, pairs are ordered by increasing genetic correlation *p*-value.

## Supplementary Note

### SN1 Empirical Parameter Estimation

Analysis with CAUSE is comprised of two main steps (see Figure S1). The first step is to estimate nuisance parameters *ρ*, which accounts for overlap in the two GWAS samples and the parameters defining the empirical joint prior distribution for *β*_*M,j*_ and *θ*_*j*_. We do this in two sub-steps. First, we select a panel of candidate covariance matrices Σ_0_, …, Σ_*K*_. Second, we fix *γ* = *η* = 0 and compute the maximum a posteriori (MAP) values of *ρ* and *π*_0_, …, *π*_*K*_.

The set of candidate covariance matrices should be large enough that a flexible joint distribution can be fit for *β*_*M,j*_ and *θ*_*j*_, but not so large that evaluating the likelihood in (7) becomes burdensome. To choose this set, we first apply the Adaptive Shrinkage (ASH) method proposed by Stephens [1] to estimate the distributions of ***β***_*M*_ and ***β***_*Y*_ separately. Briefly, given a set of summary statistics for a single study, ASH estimates a sparse unimodal distribution for the marginal effects. This distribution is flexible and parameterized as a mixture of univariate normal distributions centered at 0. ASH uses the model

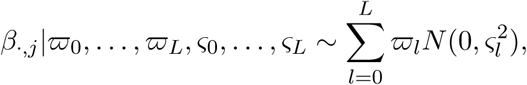

where *ς*_0_, …, *ς*_*L*_ are a fixed grid of variances with *ς*_0_ = 0. ASH estimates the mixing proportions 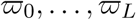 using a prior on 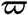 that encourages more weight to be given to 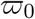, the proportion of effects equal to 0. Despite starting with a large number of candidate variances, ASH solutions tend to place most of the weight on only a few values. The resulting solution is sparse (most of the estimated effects are 0) and parsimonious (there are few components in the model with non-zero mixing proportion).

Let 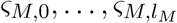 and 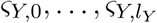 be the set of variances with non-zero weight in the ASH estimates for traits *M* and *Y* respectively. Because ASH encourages sparsity, in all cases *ς*_*M*,0_ = *ς*_*L*,0_ = 0. We construct the panel of candidate 2×2 covariance matrices by taking all pairs of these variances as diagonal elements and setting the off diagonal elements to be 0. Thus if *l*_*M*_ = 4 and *l*_*Y*_ = 3, our method produces a set of (4 + 1)(3 + 1) = 20 candidate covariance matrices.

In the second step, we fix *γ* = *η* = 0 and calculate the MAP values of *ρ* and *π*_0_, …, *π*_*K*_. We use a Dirichlet(10, 1, …, 1) prior on *π*_0_, …, *π*_*K*_ with *π*_0_ corresponding to the covariance matrix of all zeros. This prior is the same prior used by default in ASH and encourages a sparse solution, however the weights may be adjusted by the user in the CAUSE software. We use a prior on *z* = tanh^−1^(*ρ*) of *z* ∼ *N* (0, 0.25), which is a weak prior encouraging *ρ* to be close to zero. To calculate the MAP estimate, we use coordinate descent, alternating between optimization of *ρ* with ***π*** fixed and optimization of ***π*** with *ρ* fixed. As observed by Stephens [1] and others, maximization in ***π*** is a convex optimization problem that can be completed quickly. In practice, we find that convergence is usually reached within five iterations.

### SN2 Prior Distributions for *γ* and *η*

In most cases, little prior information is available about the size of the causal or confounding effect. Differences in variable scaling and covariate adjustment across GWAS may make it difficult to predict the magnitude of these effects. Fortunately, we find that CAUSE results are robust to a wide range in prior distributions for these parameters. We require that the same prior be used for *γ* and *η*. If this is not the case false positives can arise when the true shared factor effect is better represented by the prior on *γ* than the prior on *η*. We use normal prior distributions with mean 0 and variance 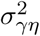 for *γ* and *η*.

To assess the robustness of CAUSE to changes in 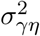, we analyze data simulated from three scenarios using a range of values for 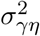. The three scenarios are 1) a setting with a causal effect 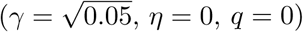, 2) a setting with no causal effect but some correlated pleiotropy 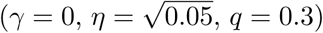, and 3) a setting with neither a casual effect or correlated pleiotropy (*γ* = 0, *η* = 0, *q* = 0). We analyze simulated data using three values of 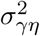. These are chosen so that 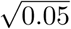, is at the 80th, 65th, and 51st quantile of the 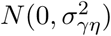 distribution, giving small, medium, and large values of *σ* _*γη*_ respectively. All simulations are conducted with sample sizes *N*_*M*_ = *N*_*Y*_ = 40, 000, the high power setting used in other simulations.

We compare *p*-values and posterior median estimates for *γ, η*, and *q* under the causal and sharing models across different values of 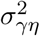. The Pearson correlation in *p*-values was greater than 0.95 for all settings and between all pairs of values of 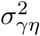. The correlation in posterior medians was higher than 0.8 for all parameters and all settings except for estimates of *q* in setting 1 under the causal model. These had a somewhat lower correlation (minimum correlation 0.51). However, in all cases the posterior median of *q* in the causal model was very low – less than 0.05. These results demonstrate that very similar inference can be obtained using a wide range of prior distributions for *γ* and *η*.

By default, 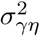 is chosen using the data. We use a set of variants with trait *M p*-value < 10^−3^ and compute 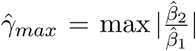. This is the largest magnitude of causal estimate that could be achieved using only one variant. We then choose 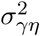 so that the prior probability that *γ* or *η* has magnitude larger than 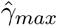 is 0.05.

### SN3 Approximating posteriors of *γ, η*, and *q*

We use an adaptive variation of a simple grid posterior approximation[2] to approximate the joint posterior distribution of *γ, η*, and *q*. To compute this approximation, we begin with initial bounds on *γ* and *η* of (−1, 1). These will be adaptively expanded as needed. The bounds on *q* are fixed at (0, 1).

The approximation proceeds as follows:

1. The domain of (*γ, η, q*) is divided into a coarse set of cubes. The approximate posterior probability of each cube is computed by approximating the likelihood within the cube as constant and equal to the likelihood at the midpoint of the cube.
2. After the first rough approximation, the bounds of *γ* and *η* are expanded so that less that 0.001 of the posterior mass falls in the cubes closest to the boundary. These bounds are then fixed.
3. The grid is then iteratively refined until no cube contains more than 1% of the posterior density. At each iteration, all cubes containing more than this are subdivided into nine smaller cubes and the posterior is re-estimated.

### SN4 Effects of LD

In this section we explore the question of whether the CAUSE model and likelihood are valid for variants in LD. For simplicity, in this section only, we assume that the GWAS for traits *M* and *Y* have no overlapping samples and have the same LD structure. CAUSE relies on two assumptions. The first is that the joint likelihood of all pairs of summary statistics can be factorized into the product of the likelihood for each variant. Variants in LD are not independent, however, by pruning variants so that the set is nearly independent, we can approximate this condition. The second assumption is that

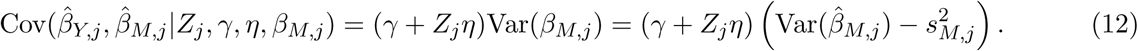

Without LD, this is a consequence of Equation (4) and the assumption that 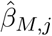 and 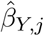 are unbiased estimates of *β*_*M,j*_ and *β*_*Y,j*_. In the presence of LD, the expectation of 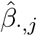 (· may be *M* or *Y*) is not *β*_*·,j*_, but a combination of *β* _*·,j*_ and a contribution from each variant in LD with variant *j*. Using results from [3],

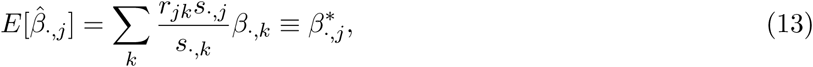

where *r*_*jk*_ is the correlation between variant *j* and variant *k*. We refer to 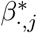 as the *LD-transformed effects*. Note that if allele frequencies are the same in the two GWAS populations then *s*_*M,j*_ = *cs*_*Y,j*_ where *c* is a constant depending on sample size. Thus

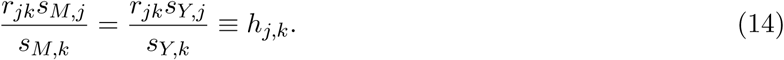

We now derive 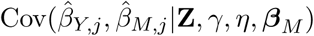 in the presence of LD, where **Z** is a vector with *j*th element equal to *Z*_*j*_ and ***β***_*M*_ is a vector with *j*th element equal to *β*_*M,j*_. We assume that direct effects are independent so Cov(*β*_*M,j*_, *β*_*M,k*_) = 0 if *j* ≠ *k* and Cov(*β*_*M,j*_, *θ*_*k*_) = 0 for all *j* and *k*. We also assume that 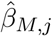 and 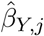 are independent conditional on 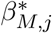 and 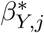. With these assumptions,

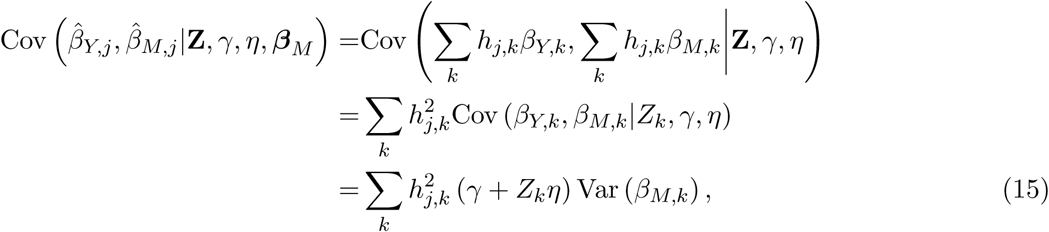

and

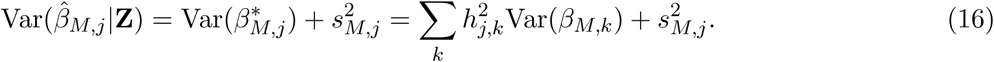

Suppose that the variant correlation structure can be decomposed into independent LD blocks and that there is at most one *M* effect variant per block. If this variant has index *k*′ then, for any variant in the block

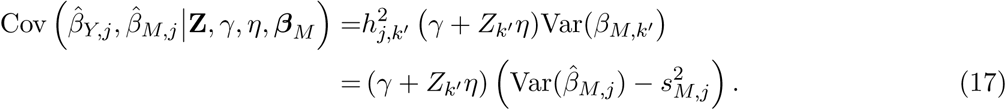

This means that, if *Z*_*k*′_ = 1, then variant *k*′ induces correlation between effect estimates for other variants in the block, even when *γ* = 0. However, if we use only one variant per LD block to estimate parameters then there is no distortion in the proportion of correlated variants. If *γ* = 0, then the proportion of variants with correlated effect estimates will be equal *q*, the proportion of true effect variants acting through *U*. In this case, CAUSE will not have an increased false positive rate but may have lower power if the variants selected to use in estimation are far from the true causal variants. To maximize power, we prune for LD prioritizing variants with low trait *M p*-values.

More generally, a block may contain multiple causal variants for *M*, with some acting on *U* (correlated pleiotropic variants) and others not. Equation 15 implies that a correlated pleiotropic variant in LD with variant *j* will induce non-zero correlation between 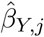 and 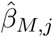 even if *γ* = 0 and *Z*_*j*_ = 0. The presence of non-shared variants in a block reduces the correlation, but will not eliminate it. If *M* is highly polygenic and *q* is large, the proportion of blocks containing at least one shared variant may be much larger than the true value of *q*. In this case, LD can create an impression that a larger propotion of variants have correlated effects, which can lead to inflated estimates of *q* and higher false positive rates using CAUSE. We assume that these settings are rare. However, our simulations are conducted under moderately dense conditions using 1000 effect variants and only 1,170 LD blocks, demonstrating that CAUSE still performs well even under somewhat unfavorable conditions.

### SN5 Connections with LCV

O’Connor and Price [4] propose an approach to identifying pairs of traits with causal relationships that uses a latent causal variable (LCV) model. This model is similar to the CAUSE model (Figure 3) with *γ* = 0. Rather than explicitly modeling both correlated pleiotropy and a causal effect, the LCV model includes only a shared factor (*U* in the CAUSE model) and estimates the “genetic causality proportion” (GCP). The GCP reflects the relative proportions of heritability of each trait that are explained by a shared factor. A causal effect (with no additional correlated pleiotropy from other sources) is equivalent to a model in which all variants act through a shared factor. In this case, the GCP is equal to 1 or −1 depending on the direction of the effect. LCV estimates the GCP and computes a test statistic testing whether GCP= 0. Models with non-zero GCP are not necessarily causal and this *p*-value is not intendd as a test of causality. O’Connor and Price [4] use an estimated GCP larger than 0.6 to suggest a possible causal relationship.

Because the LCV model is similar to the CAUSE model with *γ* = 0, we can derive an expression for GCP in terms of CAUSE parameters under this condition. The LCV model uses two parameters 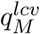 and 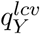, the square root of the proportions of trait *M* and *Y* heritability explained by the shared factor. Here we use the same trait *M* and *Y* notation used in our discussion of CAUSE. In terms of CAUSE parameters these are

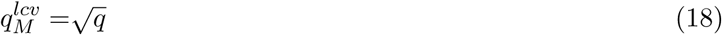

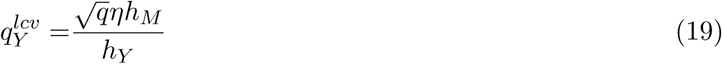

The GCP is then defined as

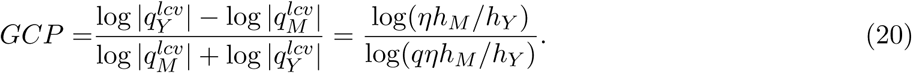

From this formula, we see that GCP is non-zero if *η* and *q* are both non-zero. For example, if the heritability of the two traits is equal, *q* = 0.3 and 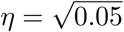 then GCP= 0.55.

### SN6 Additional Simulation Results

#### SN6.1 Effects of the prior on *q*

The choice of prior on *q* can affect the power and robustness of CAUSE. The default choice of a Beta(1, 10) prior is used in the main Results. Here we compare simulation performance using Beta(1, 2), Beta(1, 10), and Beta(1, 100) distributions. The Beta(1, 2) distribution is quite lenient, placing 25% of the prior mass on values above 0.5. The Beta(1, 100) distribution is sharply peaked close to zero, placing 95% of the prior mass below 0.03. In order for the CAUSE model to be identifiable, the prior on *q* must be asymmetric, placing more weight on values less than 0.5. Figure SN1 shows results for simulations described in the main Results (analogous to main text Figure 4). As we would expect, CAUSE with the Beta(1, 100) prior has high false positive rate when the true proportion of correlated pleiotropy is large. CAUSE with the Beta(1, 2) prior has fewer false positives for large values of *q* but also reduced power. We found the Beta(1, 10) prior to be an attractive compromise because its power is only slightly worse than power obtained using the Beta(1, 100) prior but the false positive rate is substantially improved. CAUSE shows the same performance in ROC curves for the three prior choices (Figure SN1c). One consequence of these results is that using a more permissive prior for *q* may be a better choice for higher powered studies. Results of CAUSE for pairs of GWAS traits using different priors can be seen online at the CAUSE website (see URLs).

#### SN6.2 Parameter Estimation

In the main Results, we focus on CAUSE’s performance in terms of power to detect causal effects and false discovery rate. CAUSE also produces point estimates for parameters *γ, η*, and *q* as medians of marginal posterior distributions from either the sharing or causal model. Figure SN2 compares point estimation of the causal effect, *γ*, in simulations with a causal effect and no correlated pleiotropy using CAUSE and other methods. All methods display some shrinkage. For methods besides CAUSE, shrinkage is attributed to imperfect ascertainment of instruments. In every simulated data set, there is some chance of including variants that are genome-wide significant by chance and are not tagging a causal trait *M* variant. These variants are uncorrelated with *Y* and lead to underestimation of the causal effect. As the power of the trait *M* GWAS increases, shrinkage diminishes because there are more strong *M* effect variants meeting the inclusion criteria.

Some of the shrinkage in the CAUSE estimator is also due to uncertainty about which variants are *M* effect variants. CAUSE uses all variants, and the strength of each variant’s affect on the estimate is proportionate to the strength of its association with *M*. Variants that do not tag an *M* effect variant but have strong associations with *M* by chance push the causal estimate downward. The CAUSE estimate is also shrunk by the prior on *γ* which is a wide normal distribution centered at zero. The prior on *q* does not affect the point estimate of *γ* in these experiments but does influence the width of the posterior distribution for *γ*, with narrower posteriors corresponding to stronger priors on *q* (Figure SN2(b)).

CAUSE obtains a smaller mean squared error (MSE) than Egger regression and the modal estimator for nearly every parameter value. MSE for Egger regression is not shown in Figure SN2c because it is much larger than the MSE for any other method and including it in the plot masks differences between other methods. CAUSE’s MSE is smaller or comparable to other methods for small causal effects and grows for larger effects, reflecting its shrinkage.

We find that accurately estimating *q* and *η* is more challenging than estimating of *γ*. Posterior distributions for *q* and *η* are strongly dependent and are affected by the prior distribution of *q*. In some cases, posterior distributions may be multimodal, making the posterior median a poor summary. Figure SN3 shows average posterior medians for *q* and *η* in simulations with correlated pleiotropy and no causal effect. As expected, posterior medians of *q* are shrunk more closely to zero when the prior places more weight on small values. This leads to wide credible intervals for *η* because there is little information about this parameter if *q* is small. Using the Beta(1, 2) prior for *q* we find that credible intervals for *η* are slightly narrower and that the posterior median is less shrunk relative to the true value but that credible intervals for *q* are quite wide. We are only able to obtain accurate estimates of *q* and *η* in the highest power scenarios.

#### SN6.3 Causal effects with shared factors

We explore simulation scenarios with both a causal effect and correlated pleiotropy. We focus primarily on settings with antagonistic correlated pleiotropy (correlation in the opposite direction from the causal effect), since we expect these scenarios to be the most difficult.

We first examine how a shared factor affects the power of each method. We simulate data with a causal of 0.2 and a proportion of variants, *q* = 0.1 or 0.3 acting through a shared factor with effect *η* ranging from −0.4 to 0.1. The power of CAUSE and other methods is shown in Figure SN4. A detection is only counted if the sign of the point estimate is positive. We did not find that erroneously detecting negative causal effects was a problem for any method (Egger regression detected one across all simulations and no other methods detected any). We find that CAUSE is able to maintain power when the shared factor effect has a smaller magnitude than the causal effect, but that when the shared factor effect is equal to the causal effect or larger and in the opposite direction, the power of CAUSE decreases. For larger antagonistic effects, correlation from the shared factor cancels out the correlation from the causal effect and the pattern of effect estimates becomes similar to the pattern created by a shared factor only.

**Figure SN1:**
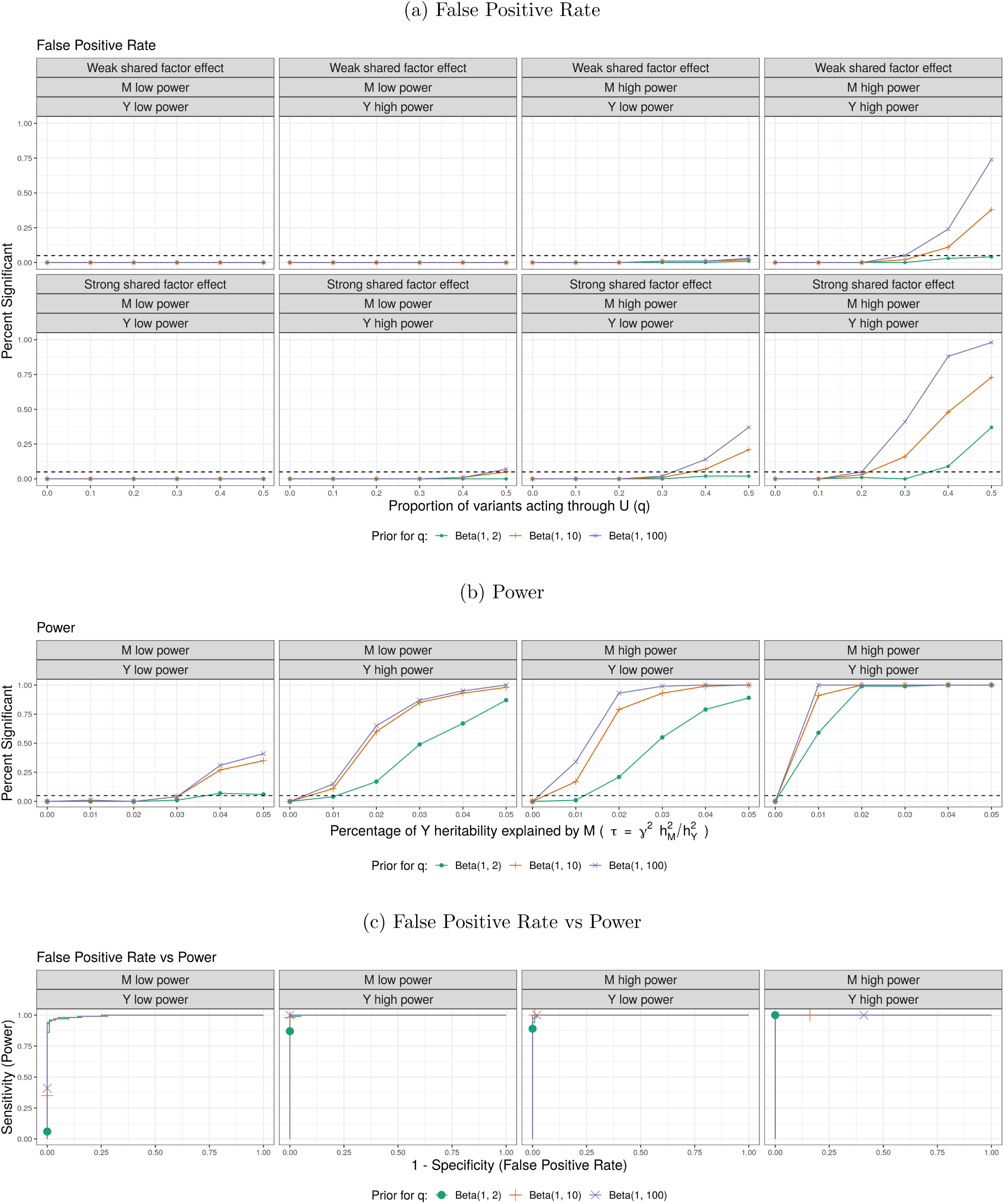
Performance of CAUSE using different priors for *q* simulated data. (a) False positive rate averaged over 100 simulated data sets in settings with no causal effect and a proportion of correlated pleiotropic variants (*q*) ranging from 0 to 50%. (b) Power averaged over 100 simulated data sets in settings with a causal effect and no shared factor. (c) Comparison of false positive-power trade-off. We compare the power when 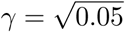 and there is no shared factor to the false positive rate when there is no causal effect, but a proportion *q* = 0.3 of variants act through a shared factor with effect 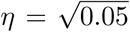 on *Y*. There are 100 simulations each in the causal and non-causal scenarios. Curves are created by varying the significance threshold. Points indicate the power and false positive rate achieved at a threshold of *p* ≤ 0.05.

**Figure SN2:**
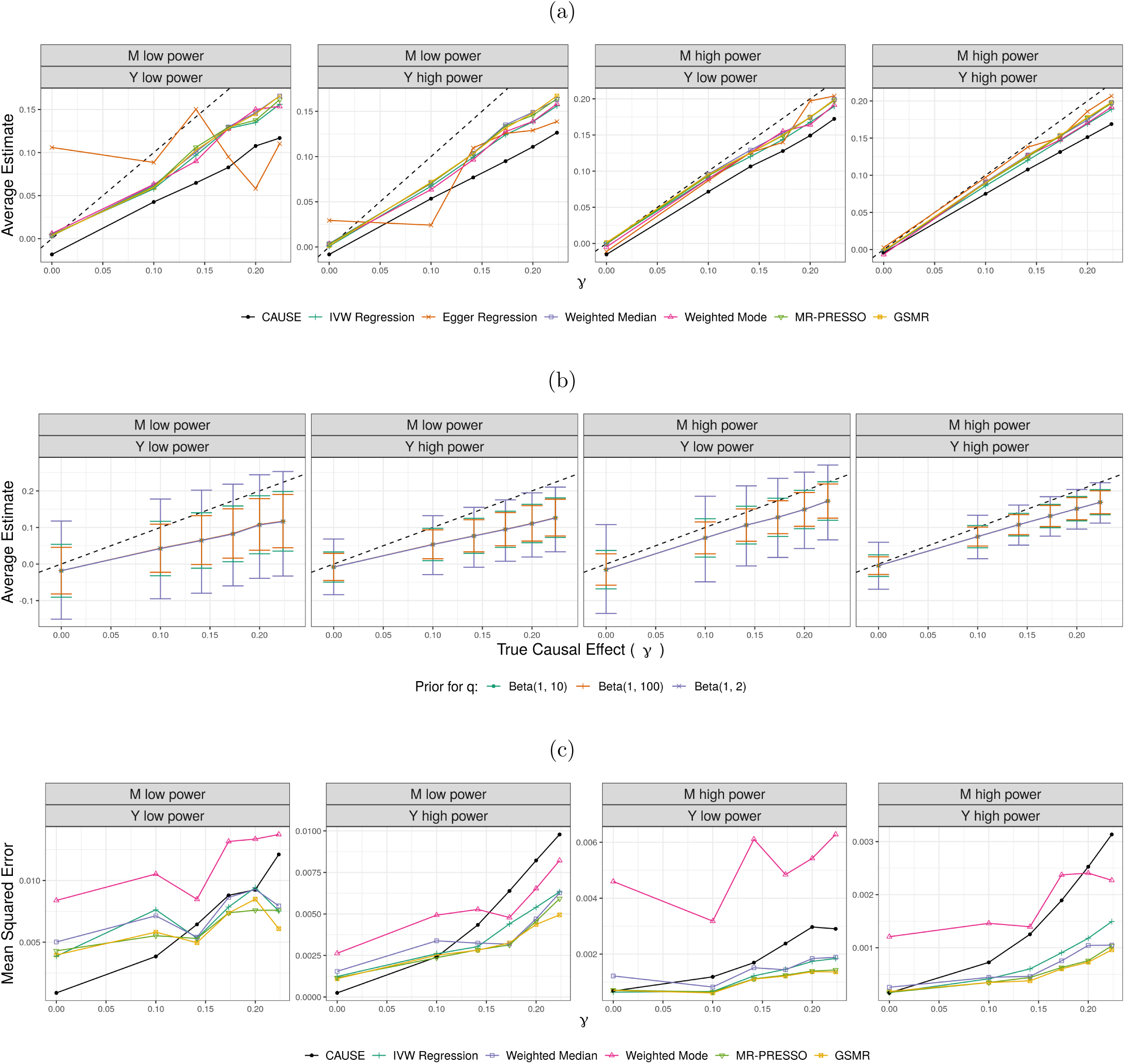
Point estimation of *γ* for simulated data with a causal effect and no correlated pleiotropy. (a) Comparing CAUSE to other MR methods, points indicate the average point estimate over 100 simulations. CAUSE is run with the default Beta(1, 10) prior for *q*. The dotted black line shows the true parameter value. (b) Comparing different priors for *q*. Error bars show average upper and lower extents of 95% credible intervals. CAUSE obtains the same point estimate using different priors but the posterior distribution is more peaked using a stronger prior on *q*. (c) Comparing mean squared error across methods. Egger regression is omitted because it obtains an MSE much larger than the other methods. CAUSE results using the default prior are shown. Note that the vertical axes in (c) are not the same across panels.

**Figure SN3:**
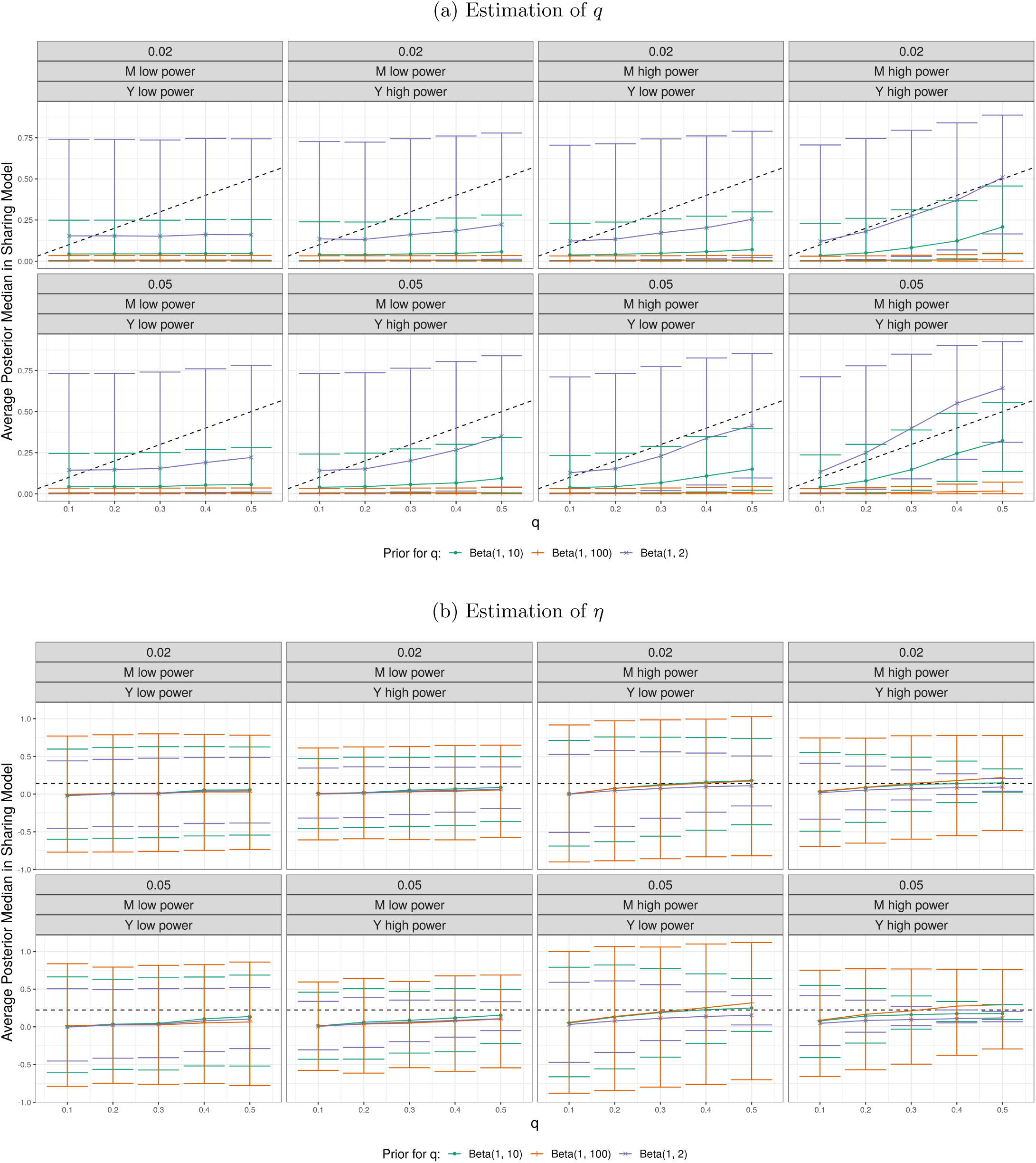
Estimation of *q* and *η* from the sharing model in simulations with no causal effect. Points show the average posterior median over 100 simulations. Upper and lower ends of vertical bars indicate the average upper and lower boundary of 95% credible intervals. Dotted lines show the true value of the parameter with estimates shown, either *q* (a) or *η* (b).

**Figure SN4:**
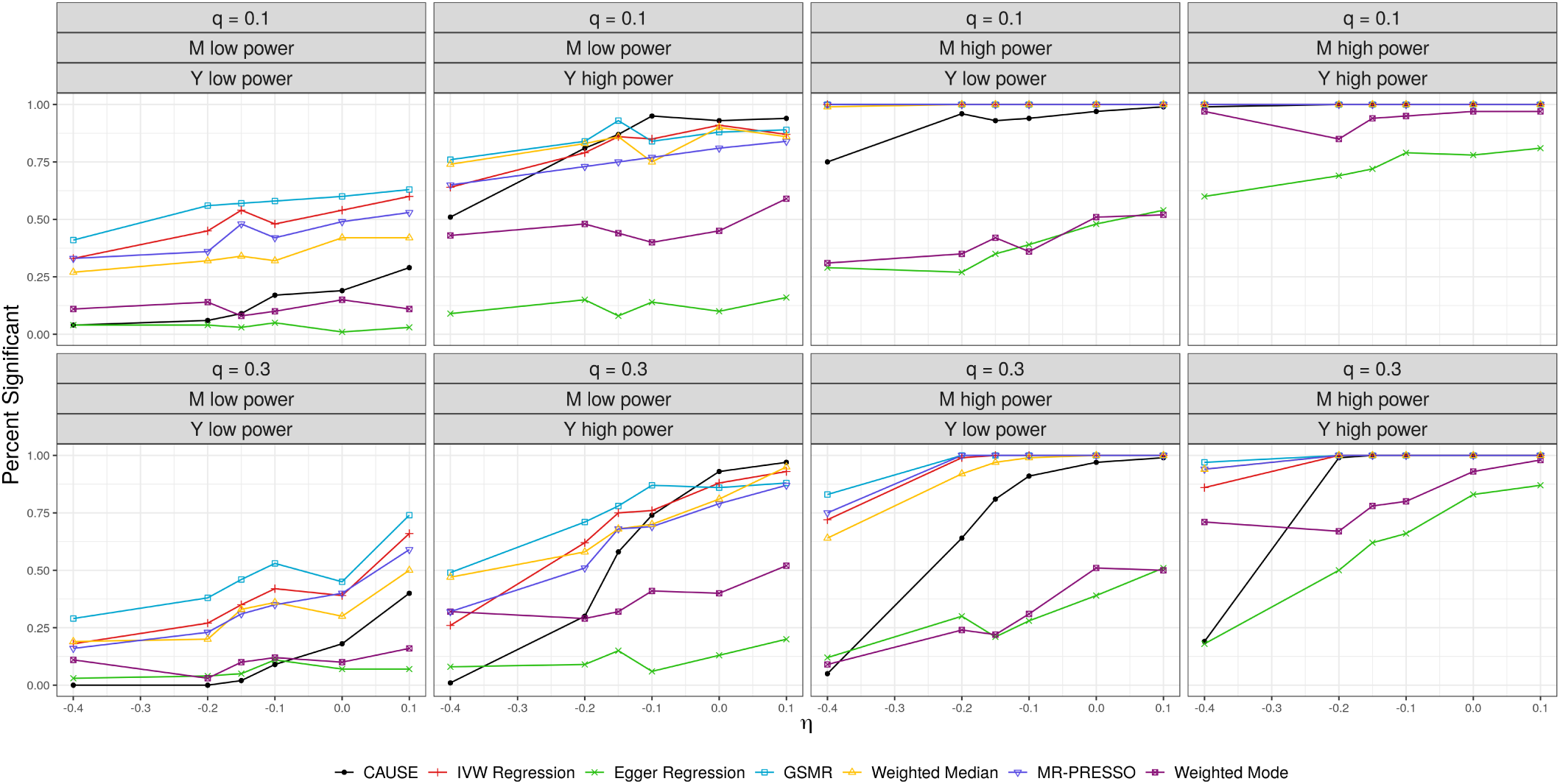
Power comparison in simulations with both a causal effect and correlated pleiotropy. In all simulations, the causal effect is 0.2. Either 10% (top row) or 30% (bottom row) of variants display correlated pleiotropy with effect size given on the horizontal axis. Points indicate the proportion on simulations detected to have a positive causal effect by each method.

Next we compare the ability of CAUSE and other methods to discriminate a scenario with a causal effect and a shared factor from one with only a shared factor. The LCV GCP estimate is included in these comparisons. In each comparison, the non-null scenario is identical to the null scenario but with the addition of a causal effect. We consider shared factors accounting for 10% and 30% of trait *M* variants and causal effects of *γ* = −1\**η* or *γ* = −2\**η*. The shared factor effect *η* is the same in all comparisons. Curves showing the trade-off between false and true positives are shown in Figure SN5. We find that, although CAUSE has lower power when there is an antagonistic shared factor, it is able to discriminate causal and non-causal scenarios at least as well as other methods.

#### SN7 Existing Information About Pairs of GWAS Traits

We provide brief summaries of literature about the relationships of the 12 risk factors and four diseases examined in the main Results. We have classified the relationship of each pair as considered causal (C), supported by literature (S), unknown (U), implausible and unsupported by literature (I), or considered non-causal (N). We classify a trait pair as considered causal if there are clinical trial or similar results supporting a causal effect and the effect is generally accepted in the literature. We classify pairs as supported by literature if there are strong molecular or physiological hypotheses supporting the effect and observational correlations. We classify pairs as implausible if there is no suggestion of a causal effect in the literature there is no genetic correlation. Effects of HDL on CAD and stroke are classified as considered to be non-causal, discussed below. All remaining trait pairs are classified as unknown or ambiguous. These include pairs that are correlated observationally but clinical trial or molecular evidence does not support a causal effect and pairs that are subject to ongoing debate. Below we summarize the literature used to make each classification, grouped by risk factor. At the end of the summary for each risk factor we list in parenthesis our classification of its relationship with CAD, stroke, T2D, and asthma in that order.

- Alcohol: Alcohol consumption is difficult to study due to its correlation with other dietary and lifestyle factors. Observational studies have suggested a protective effect of moderate alcohol consumption on cardiovascular disease including CAD and stroke [5]. This is the opposite direction of effect suggested by modal MR of alcohol on CAD, the only method to find a causal effect of alcohol on any trait at *p* < 0.05. In a clinical trial, Davies et al. [6] find moderate alcohol consumption associated with increased insulin sensitivity, and decreased triglycerides, but no effect on fasting glucose. To our knowledge, there is no evidence linking alcohol consumption and asthma. (U, U, U, I)

**Figure SN5:**
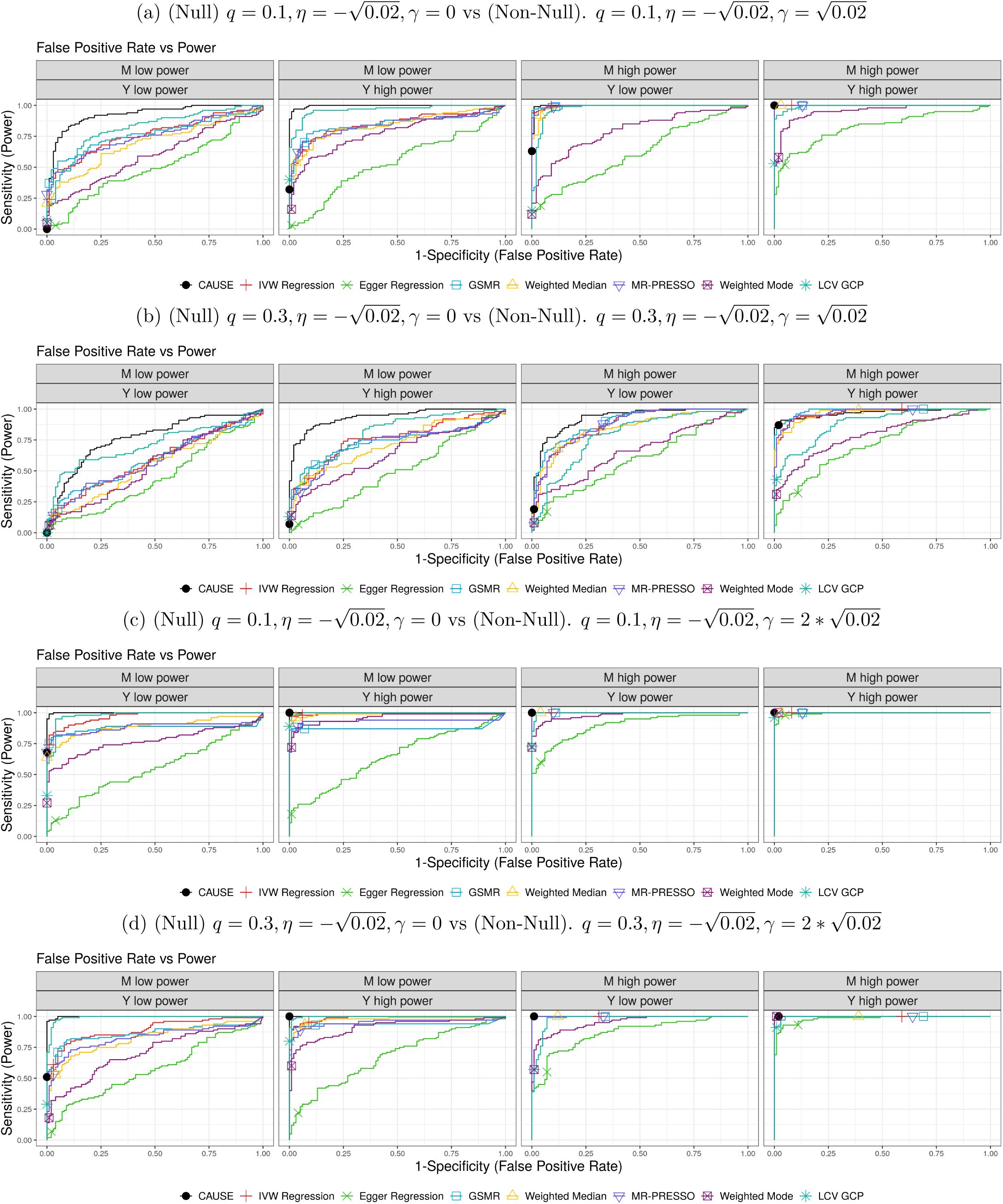
Comparison of false positive-power trade-off. There are 100 simulations each in the causal and non-causal scenarios. Curves are created by varying the significance threshold. Points indicate the power and false positive rate achieved at a threshold of *p* ≤ 0.05 or *GĈP*0.06 for LCV.
- Smoking: Smoking is a well known risk factor for cardiovascular disease (CVD), including CAD and stroke, and has been studied extensively through observational and molecular studies over the last several decades [7]. This report also concludes that smoking is a cause of T2D, and that there may be a causal relationship between smoking and asthma but this remains inconclusive. No MR methods are able to detect effects of smoking on either diabetes risk or asthma. Associations between smoking behavior and asthma risk may be confounded by effects of parental smoking behavior. (C, C, C, S)
- Blood pressure (SBP, DBP): Two recent clinical trials have studied effects of intensive blood pressure lowering in groups with high risk of CVD. The ACCORD study [8] included individuals with T2D which is associated with elevated risk of CVD. The SPRINT trial [9] included individuals with baseline systolic blood pressure above 130 mm Hg and excluding those with prior stroke. Both studies had primary composite outcomes including both myocardial infarction and stroke. ACCORD did not find a significant effect of intensive blood pressure lowering on the composite outcome but did find evidence of an effect on stroke, a secondary outcome. The SPRINT trial was stopped early due to a significantly lower rate of cardiovascular events in the intensive treatment group suggesting a causal effect. The difference between treatment groups in SPRINT was driven primarily by a decrease in heart failure and a post-hoc analysis of stroke is not significant [10]. This may be attributable to the lower stroke risk of the SPRINT population, which may reduce power to detect an effect. Although the two studies have somewhat different outcomes, we classify the relationships of blood pressure with CAD and stroke both as considered causal due to strong observational evidence and support in clinical trials. To our knowledge, no randomized trials have examined relationships between blood pressure and T2D risk or asthma risk. Because blood pressure and asthma are also not genetically correlated, we classify that relationship as implausibly causal. Blood pressure and T2D are genetically correlated so this relationship is classified as unknown. (C, C, U, I)
- Fasting Glucose: Blood glucose control and insulin sensitivity are disrupted in T2D. Impaired fasting glucose is one of the diagnostic criteria for diabetes so it would be reasonable to expect these traits to share most of their genetic variants. CAUSE finds a significant result in both directions which is consistent with this expectation. It has been suggested that elevated levels of blood glucose can contribute to T2D risk by encouraging more insulin production which eventually overtaxes the pancreas. The causal ordering between elevated is unclear plausibly reciprocal (i.e. there may be causal relationships in both directions). In a meta analysis of cohort studies, Sarwar et al. [11] find a strong correlation between T2D and CVD. However, they find that the relationship between CVD and fasting glucose is non-linear and not significant in lower ranges. It is possible that this association is mediated by diabetic disruption of lipid profiles rather than elevated blood glucose. Fasting glucose is genetically correlated with stroke and CAD but existing literature does not provide clear evidence of a causal effect so these effects are classified as unknown. Fasting glucose and asthma are not genetically correlated and have not been linked in epidemiological literature. (U, U, S, I)
- Birth Weight: Horikoshi et al. [12] study genetic effects on birth weight and genetic correlation with other traits including CAD, T2D, and asthma. They find a strong negative genetic correlation between birth weight and both CAD and T2D and no significant genetic correlation with asthma. In our study we additionally find no significant genetic correlation with stroke. Explanations for the observed genetic correlations are unknown and may result from a combination of both maternal and fetal genetics. Tyrrell et al. [13] observe a negative correlation between paternal diabetes and birth weight and a positive correlation between maternal diabetes and birth weight. This is consistent with a model in which fetal inheritance of diabetes risk alleles lowers birth weight by impeding glucose metabolism but exposure to maternal high glucose during gestation increases birth weight. Horikoshi et al. [12] suggest that the relationship between birth weight and CAD may be mediated by effects on blood pressure. If this were the case, we might also expect an effect on stroke risk, though no methods except Egger regression find evidence of this relationship in our data. (U, U, U, I)
- Height: Inverse correlations between adult height and both heart disease and stroke have been found in observational studies, with stronger and more consistent results for heart disease [14, 15]. In a Mendelian randomization study, Nüesch et al. [16] find evidence of a causal effect of height on CAD but not stroke. The authors suggest that the effect of height on CAD may be mediated by effects on lung function and lipid profiles. This study uses overlapping data and similar approaches to those used here so cannot be viewed as independent evidence. However, based on strong observational correlation and presence of a physiological explanation for a possible effect, we classify the effect of height of CAD as supported. We classify the effect of height on stroke as unknown. Height is not expected to be causally related to T2D or asthma and is not genetically correlated with either of these traits. (S, U, I, I)
- BMI and Body Fat Percentage: We use the same classifications for body fat percentage and BMI because they are closely related and, there is not always research examining the two traits separately, and we have not found qualitatively different evidence for the two traits. Overweight and obesity is often considered a risk factor for CVD. Recently, Khan et al. [17] combined data from 10 prospective longitudinal cohort studies in the Cardiovascular Disease Lifetime Risk Pooling Project to estimate lifetime risk of CVD in BMI categories. They find increased risk of CVD in the highest categories with risk increasing for groups with higher BMI. This trend was strongest for cardiac outcomes and was weaker for stroke. Observational associations are suggestive but don’t establish causality or the mechanism of association. However, because the correlations are found across studies and remain after controlling for lifestyle factors such as smoking, we categorize the relationships between BMI/body fat and CAD/stroke as supported. Obesity is typically considered a risk factor for T2D, primarily supported by consistent observational associations with higher risk in higher BMI categories [18]. Additional research has supported body fat as a stronger predictor of T2D than BMI [19]. Associations have been found between BMI and asthma in children, including in a Mendelian randomization study [20, 21]. However, the mechanism of this association is unknown and complicated by other environmental factors such as maternal BMI during gestation. It is also possible for asthma to increase risk of high BMI through decreased physical activity [22]. In a longitudinal study of a Swedish birth cohort Ekström et al. [22] observe higher BMI throughout childhood for females with persistent asthma but not for males or for children with transient asthma or asthma which onset after 4 years. Based on these findings, we categorize the relationship between BMI and body fat and asthma as unknown. (S, S, S, U)
- HDL: Low levels of HDL cholesterol are correlated with increased risk of heart disease and stroke in observational studies. However, HDL is also correlated with other known CAD risk factors including LDL cholesterol. Voight et al. [23] identify a large effect monogenic variant affecting HDL independent of LDL and triglyceride levels and demonstrate no association between this variant and CAD. Additionally, Burgess and Bowden [24] use multivariable MR to adjust for LDL cholesterol and triglyceride levels and find no effect of HDL on CAD conditional on these variables. In addition to this evidence, there are several Mendelian variants with large effects on HDL cholesterol levels that do not have clear effects on cardiovascular disease, and clinical trials of HDL raising drugs have failed to consistently show an effect. In combination, this evidence has created significant doubts that HDL cholesterol is protective for atherosclerosis as had been previously hypothesized [25]. Dyslipidemia (high LDL cholesterol, high trigylycerides, low HDL cholesterol) is often associated with T2D. This observation has been attributed to secondary effects of insulin resistance [26] but there have also been suggestions that imbalanced lipids could play a role in disease development and that particularly, HDL may be protective for pancreatic *β*-cells [27]. A Mendelian randomization study [28] using overlapping data to the data used here finds nominal associations for some tests including association between genetically higher LDL and lower risk of diabetes and weaker associations for higher HDL and lower disease risk. We note that the association between genetically predicted LDL and T2D has opposite sign from the observational association. Fall et al. [28] conclude that they have not identified consistent evidence for a causal role of circulating lipids and cite several possible confounding factors that may affect MR results. Based on this we classify the relationship of HDL with T2D as unknown. HDL cholesterol and asthma have not been linked and are not genetically correlated. (N, N, U, I)
- LDL: LDL cholesterol has been studied extensively through randomised trials of LDL lowering drugs and through MR studies of strong Mendelian LDL effect variants. Cholesterol Treatment Trialsists Collaborators [29] meta-analyze 14 statin trials and find a significant reduction in incidence of coronary heart disease and events related to vascular disease including myocardial infarction and stroke. In addition to this evidence, studies of individuals with monogenic forms of high LDL cholesterol find that these individuals are at high risk for atherosclerosis, or a build up of plaque in the arteries that can lead to heart disease and stroke and find higher incidence of CAD for individuals with monogenic high LDL cholesterol than for those without those mutations [30, 31]. Based on this evidence, we classify relationships of LDL with CAD and stroke as considered causal. Based on research summarized in the HDL section, we classify the relationship of LDL and T2D as unknown. We further note that genetic and observational studies suggest different directions of association between LDL and T2D with high LDL often observed in individuals with T2D but genetically lower LDL associated with higher risk using some methods. Statin therapy which lowers LDL has been seen to increase risk of T2D through an unknown mechanism, further complicating the picture [32]. LDL cholesterol and asthma have not been linked and are not genetically correlated. (C, C, U, I)
- Triglycerides: Like LDL cholesterol, high triglyceride levels are positively correlated with risk of heart disease and stroke. Whether this relationship is causal and if triglycerides should be a therapeutic target has been debated and there is conflicting evidence. In a summary from the American Heart Association, Miller et al. [33] note that associations with cardiovascular disease disappear after controlling for other factors including HDL and non-HDL cholesterol levels. They conclude from a range of epidemiological and molecular evidence that triglyceride levels are not directly atherogenic but are useful biomarkers for cardiovascular disease due to high correlation with levels of risk increasing particles. However, a recent meta analysis of clinical trials of lipid lowering therapies that reduce triglycerides more than they reduce LDL cholesterol does find a significant effect of triglycerides on risk of cardiovascular events [34]. This association was strongly influenced by a single outlying study and becomes non-significant when that study is removed. Two Mendelian randomization studies adjusting for LDL and HDL found a significant effect of triglycerides on CAD [35, 24]. Based on this research we classify the relationship of triglycerides and CAD as supported though we note that this issue is still debated and classify triglycerides and stroke as unknown. Based on research summarized under HDL we classify the effect of triglycerides on T2D as unknown. Triglycerides and asthma have not been linked and are not genetically correlated so we classify that relationship as implausible. (S, U, U, I)

### SN8 Discussion of selected negative results of CAUSE

In Table 1, there are several pairs of traits that are often considered to be causal and have some literature support, but CAUSE does not find a significant result. In order to understand these results more thoroughly, we examine the data and point-wise contribution to the CAUSE test statistic for these trait pairs.

#### SN8.1 Body fat percentage and CAD and T2D

Effect size plots for BF → CAD and BF → T2D are shown in Figure SN6, illustrating why CAUSE obtains a negative result. For both CAD and T2D there is significant heterogeneity in effect size correlation and slightly sub-threshold variants are not consistent with a causal effect. Causal effects of body fat percentage are found more strongly by the weighted median, weighted mode, and in the case of T2D, MR-PRESSO, methods that down-weight variants with heterogeneous or “outlying” effect size correlations.

#### SN8.2 Triglycerides and CAD

The effect of triglycerides on heart disease risk has been debated with conflicting evidence and opinions in the literature (see SN Section SN7). It is complicated by the fact that triglycerides are both genetically and observationally correlated with several CAD risk factors including LDL cholesterol. Two Mendelian randomization studies, Do et al. [35] and Burgess and Bowden [24], have tested for a causal effect of triglycerides on heart disease controlling for LDL and HDL cholesterol. Both studies find in favor of a causal effect. In the data, we see that some variants are strongly associated with triglyceride levels but not with CAD or in the opposite direction from the main trend (Figure SN7a). Some of these variants are also strongly associated with LDL cholesterol or systolic blood pressure (Figures SN7b,c), and there appears to be less heterogeneity even when variants associated with these factors are removed, though this also eliminates all variants most strongly associated with triglycerides (Figure SN7d). In order to assess whether complex pleiotropy between triglycerides, LDL, and SBP could explain CAUSE’s negative result, we ran CAUSE twice using two different subsets of variants. First using only variants with LDL *p*-value > 0.05 we obtained a *p*-value from CAUSE of 0.3 suggesting that LDL does not fully explain the heterogeneity of effect correlation. Second, using only variants with LDL *p*-value > 0.05 and SBP *p*-value > 0.05, we obtained a *p*-value from CAUSE of 0.044. These analyses are exploratory and more work needs to be done to fully understand this relationship.

**Figure SN6:**
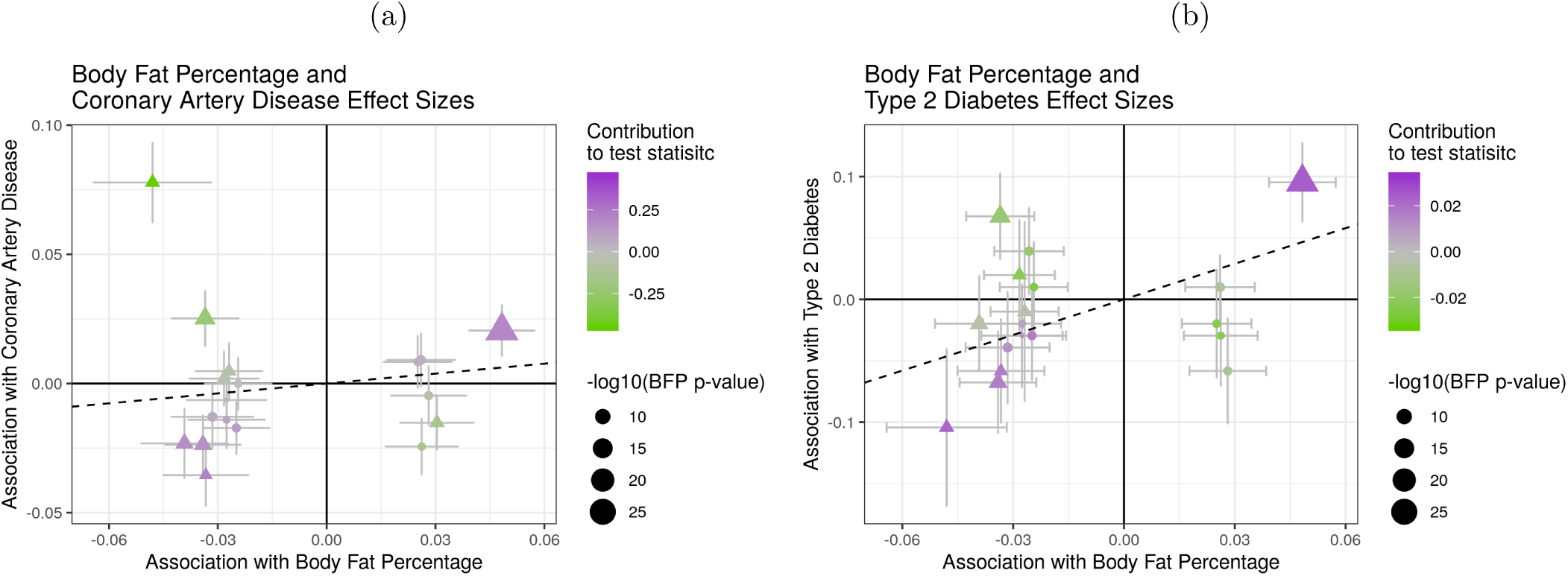
Effect size estimates and variant level contribution to CAUSE test statistics for four trait pairs. Effect estimates for trait *M* (horizontal axis) are plotted against estimates for trait *Y* (vertical axis). Error bars have length 1.96 times the standard error of the estimate. Triangles indicate variants reaching genome-wide significance for trait *M* (*p* < 5 · 10^−8^). Variants with trait *M p*-value < 5 · 10^−6^ are shown. Dotted lines show the IVW estimate obtained using only genome-wide significant variants. (a) Body fat percentage (*M*) and CAD (*Y*). (b) Body fat percentage (*M*) and T2D (*Y*).

### SN9 Results using a low powered blood pressure GWAS

We use a lower-powered GWAS for systolic and diastolic blood pressure [36] to shed light on the performance of each method. This study includes only 69,395 individuals, about 10% of the sample size of the GWAS performed by Evangelou et al. used in the main Results. We note that the Ehret et al. study is a subset of the sample used by Evangelou et al. Results using the lower powered GWAS are shown in Table SN1. Interestingly, IVW regression and Egger regression are unable to detect the effect of blood pressure on CAD using the smaller study. An explanation for this can be seen in the effect estimates for SBP and CAD plotted in Figure SN8. Only seven variants reach genome wide significance. Six have correlated effects on both traits but one highly significant variant contradicts this pattern with a negative association with SBP and a strong positive association with CAD. The weighted median, weighted mode, and MR-PRESSO all down-weight outliers through different mechanisms and are therefore able to detect the effect. CAUSE uses variants that do not reach genome-wide significance which contribute to evidence of a causal effect and models multiple sources of pleiotropy allowing it to accommodate the outlying variant and detect the effect.

**Figure SN7:**
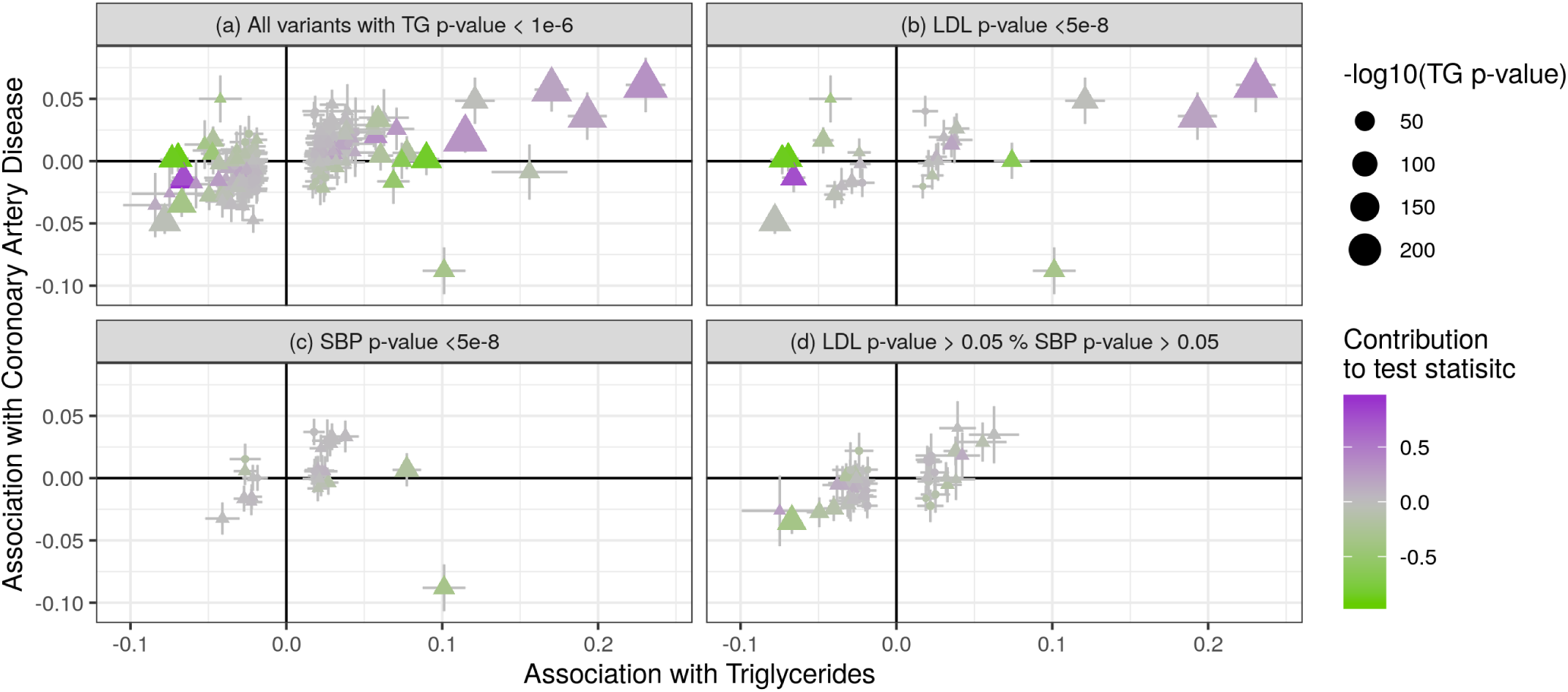
Effect size estimates and variant level contribution to CAUSE test statistics for triglycerides and coronary artery disease. Symbols and error bars as in previous plots. Dotted lines show the IVW estimate obtained using only genome-wide significant variants. (a) All variants with triglycerides *p*-value < 1 · 10^−^6. Dotted line shows MR estimate using genome-wide significant variants. (b) Only variants with LDL *p*-value < 5 · 10^−8^. (c) Only variants with SBP *p*-value < 5 · 10^−8^. (d) Only variants with LDL *p*-value > 0.05 and SBP *p*-value > 0.05.

**Figure SN8:**
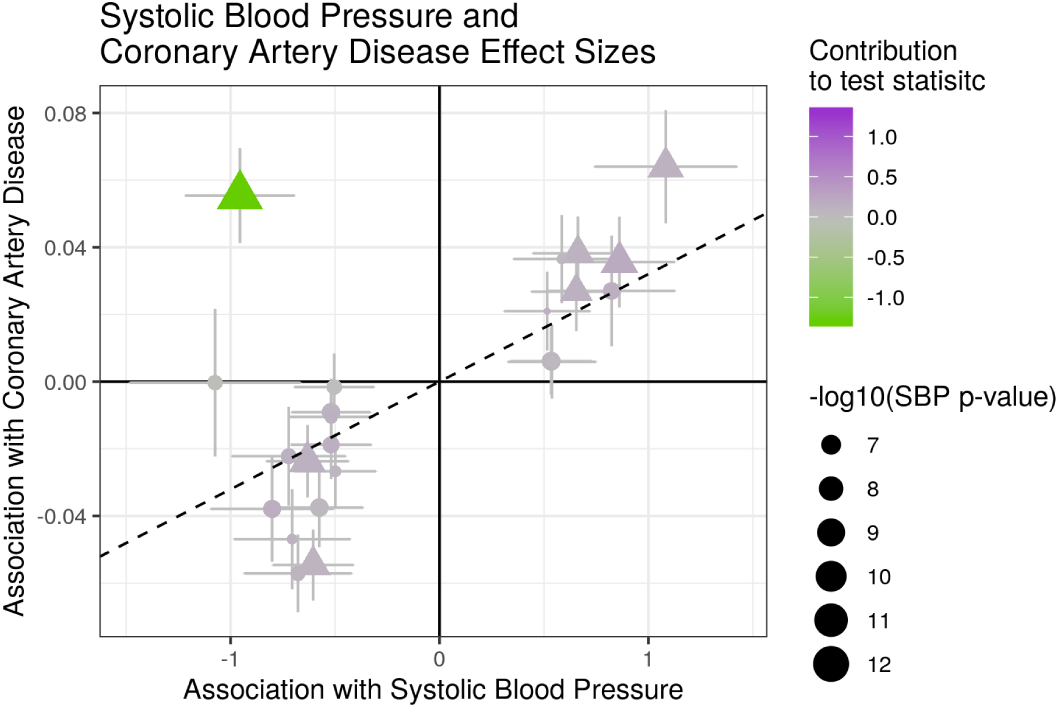
Effect size estimates and variant level contribution to CAUSE test statistics for systolic blood pressure (SBP) and coronary artery disease (CAD). Effect estimates for SBP are taken from the GWAS of Ehret et al. which has lower power than the study used in the main Results. Symbols and error bars as in previous plots. The dotted line shows the IVW estimate obtained using only genome-wide significant variants.

**Table SN1:** Summary of results using the lower powered blood pressure GWAS of Ehret et al. Columns 2-7 give the *p*-value for each MR method. Values are bold if *p* < 0.05. Arrows indicate the sign of the corresponding effect estimate. The LCV column gives the estimated GCP from LCV with *p*-value testing that GCP= 0 in parenthesis. Values are bold if estimated GCP> 0.6. The “Cause *q*” column gives the posterior median of *q* in the CAUSE sharing model and the GC column gives the genetic correlation estimated by LD score regression with *p*-value testing that genetic correlation is zero in parenthesis.

| Traits | CAUSE | IVW | Egger | Weighted Median | Weighted Mode | MR-PRESSO | LCV | CAUSE sharing $q$ | Genetic Correlation |
| --- | --- | --- | --- | --- | --- | --- | --- | --- | --- |
| SBP → CAD | **0.0021** ↑ | 0.061 ↑ | 0.43 ↓ | **$1.1 \cdot 10^{-9}$** ↑ | **$1.5 \cdot 10^{-11}$** ↑ | **0.00091** ↑ | 0.49(0.043) | 0.54 | 0.34($7.3 \cdot 10^{-15}$) |
| DBP → CAD | **0.0017** ↑ | 0.064 ↑ | 0.87 ↑ | **$4.7 \cdot 10^{-8}$** ↑ | **$1.8 \cdot 10^{-7}$** ↑ | **0.013** ↑ | 0.58(0.053) | 0.38 | 0.29($1.4 \cdot 10^{-10}$) |
| SBP → Stroke | **0.003** ↑ | **0.0053** ↑ | 0.26 ↓ | **0.00044** ↑ | **0.0037** ↑ | **0.0075** ↑ | 0.13(0.34) | 0.54 | 0.37($1.8 \cdot 10^{-7}$) |
| DBP → Stroke | **0.0014** ↑ | **$2.3 \cdot 10^{-6}$** ↑ | 0.74 ↑ | **$8.7 \cdot 10^{-5}$** ↑ | **0.0048** ↑ | **$6.9 \cdot 10^{-5}$** ↑ | 0.12(0.43) | 0.38 | 0.34($2.2 \cdot 10^{-5}$) |
| SBP → T2D | 0.22 ↑ | **0.017** ↑ | 0.29 ↑ | 0.19 ↑ | 0.32 ↑ | **0.022** ↑ | -0.28(0.35) | 0.1 | 0.18(0.0033) |
| DBP → T2D | 0.096 ↑ | 0.081 ↑ | **0.039** ↑ | 0.16 ↑ | 0.26 ↑ | 0.13 ↑ | -0.19(0.42) | 0.12 | 0.12(0.079) |
| DBP → Asthma | 1 ↓ | 0.52 ↓ | 0.33 ↑ | 0.44 ↓ | 0.59 ↓ | 0.53 ↓ | -0.06(0.77) | 0.04 | 0.06(0.37) |
| SBP → Asthma | 1 ↓ | 0.84 ↓ | 0.64 ↑ | 0.72 ↓ | 0.66 ↓ | 0.55 ↓ | -0.14(0.79) | 0.04 | 0.01(0.82) |

